# Simultaneous equations modelling of communities with interacting species networks

**DOI:** 10.1101/2021.03.07.434253

**Authors:** Miguel Porto, Pedro Beja

**Affiliations:** CIBIO/InBio – Centro de Investigação em Biodiversidade e Recursos Genéticos, Universidade do Porto, Campus de Vairão, 4485-661 Vairão, Portugal; CIBIO/InBio – Centro de Investigação em Biodiversidade e Recursos Genéticos, Instituto Superior de Agronomia, Universidade de Lisboa, Tapada da Ajuda, 1349-017 Lisboa, Portugal

**Author notes:** corresponding author: Miguel Porto. **Statement of authorship:** Both authors contributed significantly to the manuscript. MP developed the statistical methods and the software package, analysed the data and collected the data of the case study.

**Keywords:** Assembly processes, biotic interactions, community modelling, environmental filters, latent variables, interaction networks, multivariate occurrence data, network topology selection, joint species distribution model

## Abstract

To understand community assembly, ecologists have long sought to extract the signal of biotic interactions from species co-occurrence patterns. These efforts face multiple difficulties such as confounding environmental effects, confounding indirect interactions between multiple species and asymmetry of interactions. To address these problems, we propose Simultaneous Community Equations Modelling (SCEM) as a framework to explicitly account for asymmetric interaction networks in community models. SCEM uses a system of equations to model the occurrence of each species as a function of measured and unmeasured (latent) environmental predictors, and the occurrence of potentially all the other species in the community. Biotic interactions most supported by the data are identified using heuristic optimization of a parsimony criterion, implemented as a Genetic Algorithm. Extensive simulations show that SCEM can recover interaction network topologies in virtual communities. We present a software to implement SCEM and illustrate its application with a case study.

## Introduction

There is general agreement that the assembly of local species communities is shaped by the interplay of multiple drivers, including the species pool, neutral processes, speciation, dispersal, abiotic environmental factors and biotic interactions (Götzenberger *et al*. 2012). To gain quantitative insights on the relative contributions of these drivers, ecologists often use empirical, model-based approaches that combine observational data and sophisticated statistical frameworks. These frameworks have been increasingly used to explain and predict variation in the identities, numbers and abundances of species over a range of spatial and temporal scales (Norberg *et al*. 2019). However, they are not without problems, having limitations that may hinder their ability to provide a mechanistic understanding of community assembly processes (Dormann *et al.* 2018, Blanchet *et al.* 2020).

An important challenge in community modelling is to detect the signal of biotic interactions in observational data (Dormann *et al.* 2018; Blanchet *et al.* 2020). Although interactions between species are expected to affect their joint spatial patterns of occurrence (co-occurrence), spatial associations between species are a poor proxy for biotic interactions because they reflect the mixed result of multiple processes also affecting co-occurrence (Nieto-Lugilde *et al.* 2018; Caradima *et al.* 2019). For instance, spurious species associations may be inferred from shared responses to unmeasured drivers (missing predictors) (Warton *et al*. 2015), while the signal of pairwise interactions may be confounded by other interactions in a community (Cazelles *et al*. 2016). Because of this, serious concerns have been raised to the inference of ecological interactions from co-occurrence data (Dormann *et al*. 2018; Blanchet *et al*. 2020).

In their thorough review, Blanchet *et al*. (2020) recently showed that these problems (apart from those arising from inadequate sampling or data) are mainly rooted in a central issue: seeking the signal of interactions using symmetric species associations, which measure the probability of co-occurrence without accounting for the possible interactions with other species and for other processes that cause species associations. Not accounting for all the other interactions can significantly lead to erroneous results, either false positives or false negatives. Moreover, interactions are not symmetric, hence, any model that attempts to infer a species interaction from the symmetric association of two species is flawed by construction (Box 1; Cazelles *et al*. 2016; Blanchet *et al*. 2020). Yet, despite the multiple arguments against inferring interaction from co-occurrences (Dormann *et al*. 2018; Zurell *et al*. 2018; Blanchet *et al*. 2020), this is what most current models still do, similarly to other non-model-based techniques: they base the inference on some kind of quantification of (residual) species associations (Gotelli & Ulrich 2010; Warton *et al.* 2015; Ovaskainen *et al.* 2017a; Niku *et al.* 2019; Pichler & Hartig 2020). This has, in fact, become a common practice in the ecological literature (D’Amen *et al*. 2018; Abrego *et al*. 2020), encouraged by the large amounts of presence-absence data available nowadays and the rich toolset of modern statistical methods tailored to analyse this kind of data (Warton *et al*. 2015; Ovaskainen *et al*. 2016, 2017a; Nieto-Lugilde *et al*. 2018; Niku *et al*. 2019; Tikhonov *et al*. 2020a, b; Tobler *et al*. 2019; Pichler & Hartig 2020).

A model aiming to incorporate the effects of biotic interactions must, at least, account for their asymmetric nature and control for all the other interactions in the community, which is not possible using association matrices or other forms of co-occurrence analysis. A better approach is to include the effects of each species on each of the others explicitly in the model, taking the so-called interactor-as-predictor approach (Dormann *et al*. 2018). This idea is not new (see, e.g., Wisz *et al*. 2013; Harris 2016; Sander *et al*. 2017), but it has hardly been implemented in practice, because it becomes highly demanding in terms of processing power as the number of species increases (Pichler & Hartig 2020). The most sophisticated attempts to couple an explicit interaction network model to a classical Species Distribution Model, which use Markov networks (Harris 2016) or Bayesian networks (Staniczenko *et al.* 2017), still have important limitations. As Blanchet *et al*. (2020) pointed out, the full topology of the network needs to be known beforehand, which is generally impractical, as that is usually what needs to be discovered. Gaussian copula graphical models (Popovic *et al.* 2019) allow estimating the network topology but, similarly to the Markov network approaches, use undirected graphs, i.e. model symmetric species associations, hence share similar problems to co-occurrence based methods (Blanchet *et al*. 2020). On top of this, all these network-explicit models are unable to estimate unmeasured predictors along with biotic interactions (Harris 2016; Staniczenko *et al*. 2017; Popovic *et al*. 2019), because they are not joint models in essence. As such, they suffer from the same core limitation of correlation-based models: potentially confounding unmeasured environmental drivers with species interactions.

To address these problems, we describe in this paper a new community modelling framework that incorporates an explicit model for asymmetric species interactions *combined* with a model for the abiotic component, which includes unmeasured environmental predictors (Fig. 1). We formulate it as a Simultaneous Equations Model (Wisz *et al.* 2013): a system of interconnected equations where each species is included as a predictor of each of the other species, while incorporating also environmental predictors and latent variables. Simultaneous Equations Models have been extensively used in Econometrics for problems involving feedback loops between quantities, like supply and demand, and for which we seek an estimate of the reciprocal effects between entities, at some kind of equilibrium state (Rothenberg 1990; Baltagi 2008). This is actually homologous to the functioning of biological communities with interacting species (excluding some extreme cases of rapidly changing communities), where all species potentially interact, their abundances may fluctuate in response to one another, but the long term trend is steady (Soliveres *et al*. 2015; Levine *et al*. 2017). The proposed formulation, therefore, allows estimating species biotic interaction networks from occurrence/abundance data, while controlling for the confounding effects of shared responses to measured and unmeasured environmental drivers and for the effects of all other interactions in the community. Inference of interactions is made directly from the estimation of asymmetric interaction coefficients instead of from shared responses to latent variables or from residual species associations, thus stepping over multiple problems inherent to the inference based on co-occurrence probabilities (Blanchet *et al*. 2020; see also Box 1).

**Figure 1:**
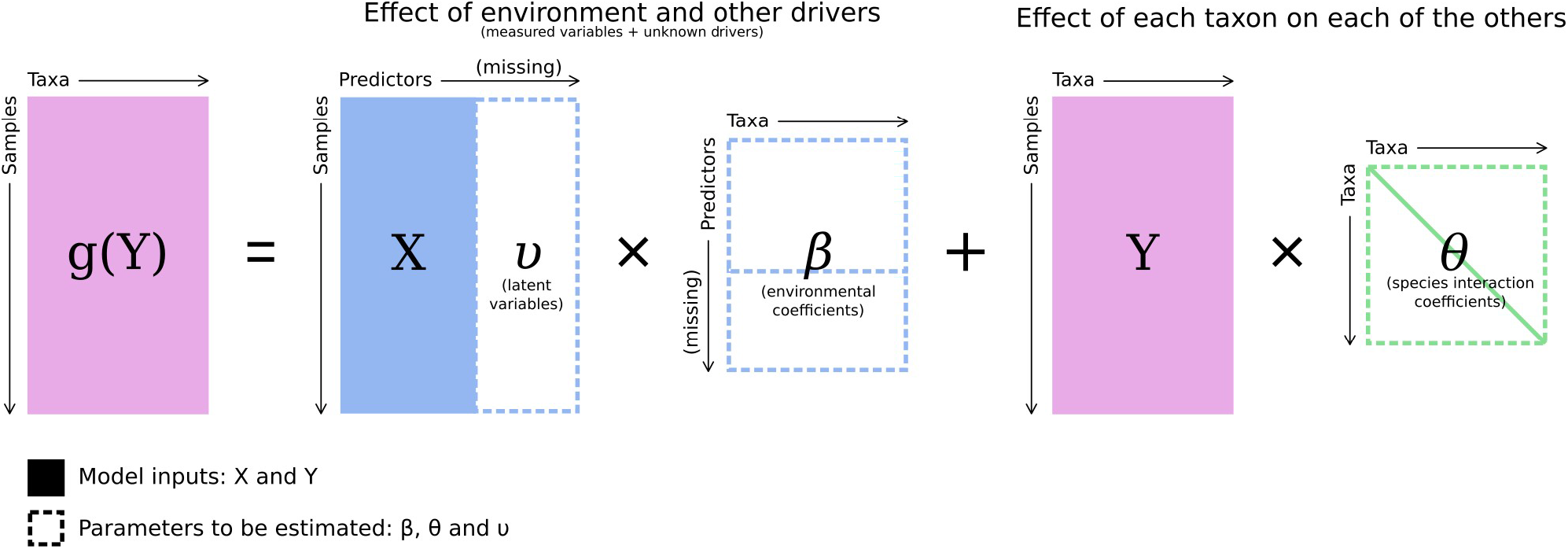
The Linear Predictor of the SCEM, in matricial form. *X* is the environmental predictor matrix, which includes missing predictors to be estimated, *Y* is the species presence-absence or abundance matrix in each site, *g* is the inverse link function. Note that the θ matrix is asymmetric and has a zero diagonal.

Fitting is implemented through Penalized Maximum Likelihood, to overcome problems of overfitting and multicollinearity in a model involving a system with potentially hundreds of equations and variables. Because the model can produce thousands of species interaction coefficients for all the species pairs in the community, we further propose a network topology selection algorithm based on heuristic optimization of a AIC-like parsimony criterion, implemented as a Genetic Algorithm (Calcagno 2020). Using simulations based on occurrence data in virtual communities and taking realistic assumptions, we show that our framework is able to a) recover, to a very reasonable extent, the true interaction network topology and interaction strengths; b) disentangle the effects of species interactions from those of shared responses to measured and unmeasured predictors; c) disentangle direct species effects from those mediated by other species, such as in cases where the true interaction network has complex effect chains with many indirect effects present; and d) recover the direction of the interactions to a reasonable extent. To facilitate the adoption of our framework by ecologists, we illustrate its application using a real example, and briefly present the R package ‘eicm’ to implement it in practice (Porto & Beja 2020).

## The Simultaneous Community Equations Model (SCEM)

### Model definition

The Linear Predictor (LP) of a SCEM in its simplest formulation (Fig. 1, Appendix S1 in Supporting Information) is a simple extension of a system of independent, single species models, where each species’ LP includes all the other species as predictors, along with all the environmental predictors, of which some, or all, can be unknown. It takes the form η = X’β + Yθ, with X’ = [X υ] and θ_ii_ = 0 (Appendix S1). If Y is a presence-absence matrix (Y_ij_∈ {0, 1}), the log-probability of observing a given species × sample matrix Y given the model parameters β and θ, and the environmental predictors X’ (i.e. the log-likelihood function for a given model parameterization) is given by

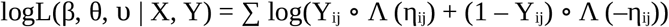

with Λ the logistic function and ∘ standing for the element-wise matrix product. Y is the species × sample presence/absence matrix (dimensions: n × m) and X is the site × environmental predictor matrix (dimensions: m × k). This formula can be read as the sum of the log-probability of each species presence or absence, given the environmental predictors and the presence/absence pattern of all the remaining species at that site. The environmental coefficients to be estimated are organized in the k × n matrix β, but in contrast with other methods (Ovaskainen & Soininnen 2011; Pollock *et al*. 2014), no assumptions are required regarding any common distribution upon these coefficients. The model is, therefore, non-hierarchical (sensu Ovaskainen & Soininnen 2011).

The species interaction coefficients are organized in the n × n matrix θ, also assumed to be unstructured, representing the effect of species j (columns) on species i (rows). This matrix has n × (n – 1) coefficients, because interactions are not assumed to be symmetric and diagonal elements are obviously removed. The unknown environmental predictors, or latent variables, comprise the matrix υ, which has m × u values to be estimated. In contrast with other methods (e.g., Hui *et al*. 2015; Ovaskainen *et al*. 2016; Niku *et al*. 2017), we also don’t assume any distribution upon the latent variables, nor their independence. For notational convenience, it is concatenated with the known environmental predictors X (if any). The number of model parameters to be estimated under a full interaction network is then given by p = (k + u + 1) × n + n × (n – 1) + m × u, where n is the number of species, k is the number of measured environmental predictors, u is the number of unknown environmental predictors, m is the number of sampling sites and the + 1 stands for the species-level intercept.

This innovative model architecture allows unmixing the two major confounding sources of co-variation, i.e., shared responses to unmeasured covariates and species interactions, which is a novel feature that is not possible in any of the existing models for snapshot data, either network-explicit or association-based (Ovaskainen *et al*. 2016; Niku *et al*. 2017; Staniczenko *et al.* 2017; Pichler & Hartig 2020). This becomes possible because, in a multi-species setting, the signal of shared responses to environment is very likely to span across multiple species, thus emerging in the top latent factors because of its wide community-level effect. Contrariwise, the signal of species-specific interactions has only a restricted effect that is not likely to be captured in the top latent factors (Pichler & Hartig 2020), therefore, emerging as individual interaction coefficients in our model but likely to be either missed in the current approaches relying on latent factors (Ovaskainen *et al*. 2016; D’Amen *et al*. 2018; Wilkinson *et al*. 2019) or diluted within the residual association matrix, in the latent factor free approaches (Pichler & Hartig 2020).

### Parameter estimation

Model parameters are estimated via penalized Maximum Likelihood, while the respective confidence intervals are estimated via likelihood profiling. No approximations, simplifications or assumptions are needed by the estimation algorithm. All the parameters, including the values of the latent variables at each site, are estimated numerically without assumptions of their distribution or joint distribution. When u = 0 (no missing predictors), there are no shared parameters between species, hence the likelihood surface is unimodal along each independent parameter. This allows the usual gradient-based optimizers for ML estimation (with finite-difference gradient approximation) to be used, which guarantee convergence at the global maximum. When estimating the model with unknown predictors (i.e. u > 0), both the X’ and β matrix of the term X’β in the LP have unknowns, which causes the likelihood surface to become multimodal. In particular, when u > 1, the likelihood surface has flat regions, as there are infinite linear combinations (rotations) of the estimated latent variables and corresponding species coefficients yielding the same value of the LP (cf. proof in Appendix S1), thus, the same likelihood. This rotational invariance of the latent variables does not preclude the use of gradient-based optimizers for estimating species interactions, as long as we impose some constraint in the model to cancel it out (Warton *et al*. 2015). In our case, we fix the latent variables at one specific rotation prior to estimating interactions.

An important issue that results from the very high model complexity is overfitting, particularly when latent variables are estimated, because of the large number of parameters that are included in the model (Bell & Schlaepfer 2016). Another problem is multicollinearity, especially when complex species interaction networks are estimated, because species are themselves included as predictors of other species (Pollock *et al*. 2014). If not controlled, these problems would cause many inflated, unstable and boundary estimates (Hoerl & Kennard 1970; Heinze & Schemper 2002; Hutchinson *et al*. 2015). This is the reason why we use penalized likelihood, with a penalty term akin to that of ridge regression (λ ∑ β^2^) or of LASSO regression (λ ∑ |β|). The penalty plays two roles: a) ensuring the estimates are finite, which is essential when estimating missing predictors; and b) shrinking spurious species interaction estimates to zero (Popovic *et al*. 2019). Since the model has two distinct parameter matrices to be estimated (environmental and biotic interactions), we separate the penalty term into two components λ_β_ (penalty applied to the environmental coefficients and latent variables) and λ_θ_ (penalty applied to the species interaction coefficients), because these matrices are quite different in terms of size and nature, often requiring, in practice, different values for λ. The objective function of the optimization algorithm is then Q = logL(β, θ, υ | X, Y) – λ_β_ P_β_ – λ_θ_ P_θ_, where logL is the log-likelihood function and P the penalties, which may either be P_β_ = ∑ υ^2^ + ∑ β’^2^ or P_β_ = ∑ |υ| + ∑ |β’| and P_θ_ = ∑ θ2 or P_θ_ = ∑ |θ|, where β’ is the β matrix without the species-level intercepts.

### Interaction network topology

Although it is straightforward to estimate all the model parameters under the assumption of a full interaction network, such full estimation is hardly useful for ecological inference due to the large amount of spurious interaction terms, reflecting overfitting. To prune spurious species interactions, hence having a better approximation to the “true” biotic interaction network, we developed a variable selection procedure on network topology that penalizes species interaction terms similarly to the AIC-based model selection procedures. However, since it is intractable to make a full model selection table of all possible network topologies, we use a heuristic optimization that aims to find an approximation of the most plausible subset of interactions without calculating all possible models. For this, we use a Genetic Algorithm (as in Calcagno 2020) in which the decision variables control the inclusion/exclusion and direction of any given species interaction. At each algorithm iteration, the candidate models, each corresponding to a particular directed network topology, are fitted, and their fitness is calculated as the penalized log-likelihood of the model plus a penalty term for the number of interaction terms included, weighted by a factor λn. This factor controls the trade-off between Type I and Type II errors, in respect to the detection of interactions. To decrease complexity of the network selection, and thus increase the chance of finding the most parsimonious interaction network, the selection can be conducted on a reduced network model. First, the user may provide a list of forbidden and/or allowed interactions, which can be straightforward in many situations (Morales-Castilla *et al*. 2015). In addition, the model can exclude interactions that depart from rare species (with a user-specified threshold), because these can hardly be estimated anyway. Moreover, complexity can be further alleviated by excluding weak interactions from a preliminary full-network estimation, based on a user-specified threshold, θ_0_.

### Model implementation

Our model is implemented in the R package ‘eicm’ (Porto & Beja 2020), which at present can be used only with presence/absence data. The basic information required is a community matrix describing species presence/absence (0/1) at sampling sites. An environmental matrix with abiotic variables measured at sampling sites can also be used, though this is not compulsory because environmental variation can be described through latent variables. Given the model complexity, fitting should follow the staged workflow described in Appendix S1, which is implemented in the main function of ‘eicm’ and has been extensively tested: 1) estimate and fix latent variables; 2) prune the network model based on preliminary interaction estimates; 3) select network topology departing from the reduced model. This workflow requires setting five hyperparameters that control the complexity of the model and define the penalties used: λ_β_, λ_θ_, λ_n_, θ_0_ and λ_υ_ (see Appendix S1 for definition). The package is prepared to run the computer-intensive tasks in parallel, in particular the selection stage, therefore taking advantage of multi-core computing to have linear gains in speed.

### Case study

We illustrate the application of our framework using a real dataset involving the presence/absence of 300 plant species at 109 sampling sites in a Mediterranean region (Appendix S2). We also used an environmental matrix including three climatic and three topographical variables (centered and scaled to unit variance) expected to affect community composition in the study area. To control for environmental or spatial gradients that might not be captured by the measured variables, we considered six latent variables. This number was chosen as a compromise following preliminary trials suggesting that a lower number would still not capture all important community gradients, resulting in many unlikely species interactions from an ecological point of view, and a higher number would become too conservative resulting in nearly no recovered interactions. Given the very high number of potential (asymmetric) interactions, the analysis was restricted to interactions among woody plant species (i.e., trees and shrubs), and between these and species with other life forms, thereby avoiding an exceedingly complex interaction network model. In principle, other restriction criteria might also be used, as for instance assessing only potential interactions among herbaceous species. Additionally, we excluded interactions of rare species (< 10 occurrences) on other species, though we retained the reverse possibility (i.e., other species affecting rare species). The threshold of 10 was deemed sufficient for controlling model complexity while minimizing the loss of important biotic signals. The resulting restricted network comprised 7200 possible directional interactions to be estimated. The SCEM model was fitted with the following hyperparameterization: λ_β_ = 5, λ_θ_ = 2, λ_n_ = 1.5, θ_0_ = 0.5, λ_υ_ = 1. These values were based on the results obtained with simulated data that performed best in the trade-off of eliminating spurious interactions while keeping true positives (see model testing results), but were adjusted to be less conservative in the interaction penalties (λ_θ_ and λ_n_).

The fitted SCEM model supported the view that our community was very strongly structured by environmental drivers, but also that there were potentially important biotic effects. Regarding the environment component, the model highlighted the strong influence of the six measured environmental variables, but also that a more thorough understanding of community assembly processes required the specification of latent variables. In fact, the latent variables captured an important component of unexplained environmental variation, probably reflecting local scale abiotic factors that could not be adequately captured by our variables measured at relatively coarse scales (Appendix S2, Fig. S1 in Supporting Information). Regarding the biotic component, the model supported the occurrence of 11 meaningful interactions that were retained after the selection procedure (Fig. 2), indicating that the probability of a species to be present in the community is affected (positively or negatively) by the presence of another species. For instance, there was a negative effect of *Quercus suber* (cork oak) on the tiny understorey fern *Anogramma leptophylla*, even after accounting for the opposite responses of both species to the latent variable #1 (Fig. 2). This means that their mutually exclusive co-distribution pattern could not be sufficiently explained by opposite responses to a latent gradient, requiring a negative species interaction to explain the pattern. On the other side, a positive effect of *Rubus ulmifolius* (bramble) over the woodland grass *Brachypodium sylvaticum* was identified, even though these two species responded in opposite direction to latent variable #1, which might be depicting, for example, a sheltering effect of the bramble over this shade-loving grass.

**Figure 2:**
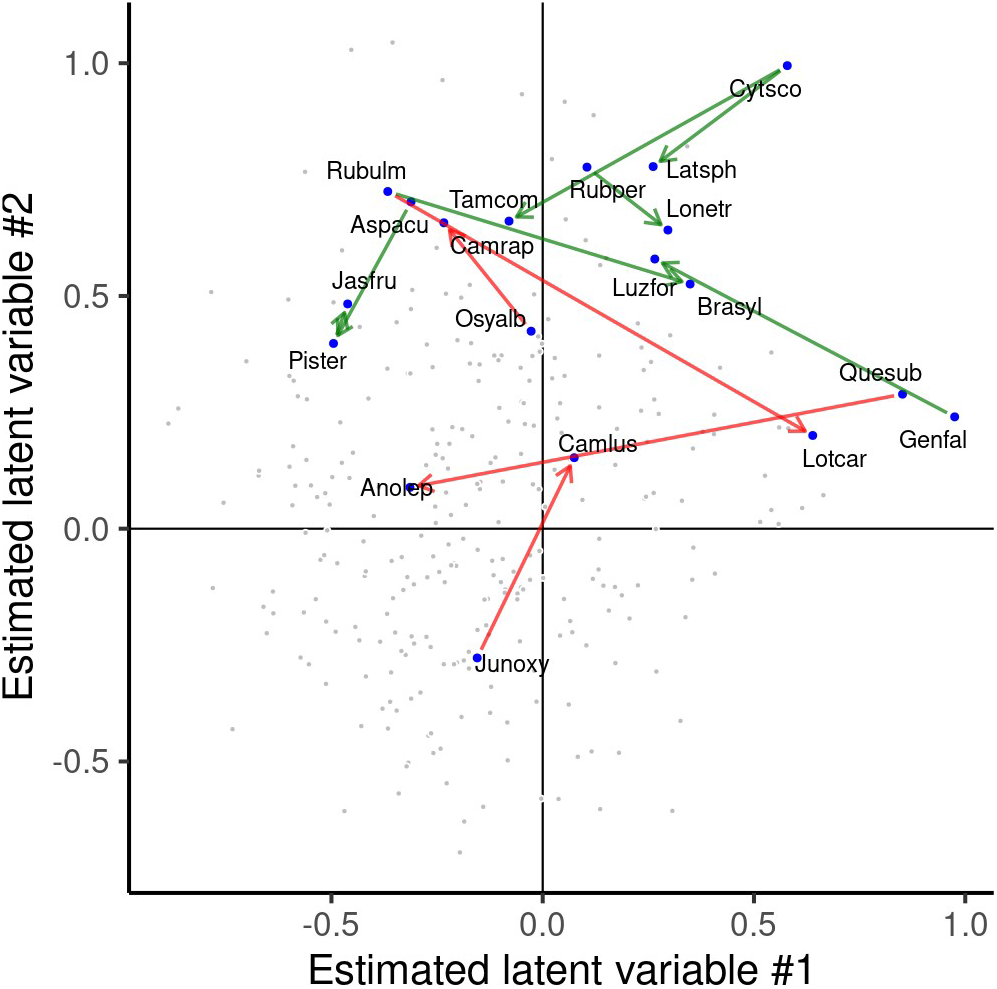
Estimated species interactions (arrows) plotted in the latent variable space of the first two estimated latent variables, of a SCEM fit to real data. The arrows depict the effects of one species on the other (arrow direction) that were not captured by the latent variables, thus illustrating the ability of the model to separate the effects of shared unmeasured predictors (axes) from the effects of species interactions. Only the estimated species interactions that were selected by the selection algorithm are plotted. Green arrows correspond to positive effects, red arrows to negative effects. Gray dots are the species for which no interaction was detected. Anolep: *Anogramma leptophylla*, Aspacu: *Asparagus acutifolius*, Brasyl: *Brachypodium sylvaticum*, Camlus: *Campanula lusitanica*, Cytsco: *Cytisus scoparius*, Genfal: *Genista falcata*, Jasfru: *Jasminum fruticans*, Junoxy: *Juniperus oxycedrus*, Latsph: *Lathyrus sphaericus*, Lonetr: *Lonicera etrusca*, Lotcar: *Lotus corniculatus* subsp. *carpetanus*, Luzfor: *Luzula forsteri*, Osyalb: *Osyris alba*, Pister: *Pistacia terebinthus*, Quesub: *Quercus suber*, Rubulm: *Rubus ulmifolius*, Rubper: *Rubia peregrina*, Tamcom: *Tamus communis*.

## Model testing with simulated data

To assess the accuracy of the model in recovering the true species interaction network from occurrence data in the presence of confounding factors, we conducted a battery of trials with simulated data for 64 virtual species under different model hyperparameterizations (details in Appendix S3). Each simulation trial included a) the generation of a presence/absence community matrix with a known environmental and species interaction model; b) fitting and selecting the model to that matrix with a given hyperparameterization, dropping all environmental predictors used in data generation; and c) assessing the accuracy of the estimates by comparing with the true model the recovered species interactions and network topology. The same protocol was implemented for virtual communities with 128 species, though here we only tested a few hyperparameterizations. Care was taken to ensure maximum realism in the simulation setting and avoid tautological results.

## Results

Results for simulated communities with 64 species and 50 true interactions with two unmeasured predictors showed that the fitted model, after network selection, was able to correctly identify between 50% and 70% of the true interactions whose coefficient was 0.5, or higher, in absolute value (|θ| >= 0.5), while selecting a small number of false positives (in the best cases, lower than 5), from a universe of 2016 possible interactions that could have been selected (Fig. 3). This trade-off between true positives and false positives depended on the hyperparameterization of the model, with the penalty for the interaction terms (λ_θ_) and for the number of interactions included (λ_n_) playing an important role in the trade-off (Fig. 3).

**Figure 3:**
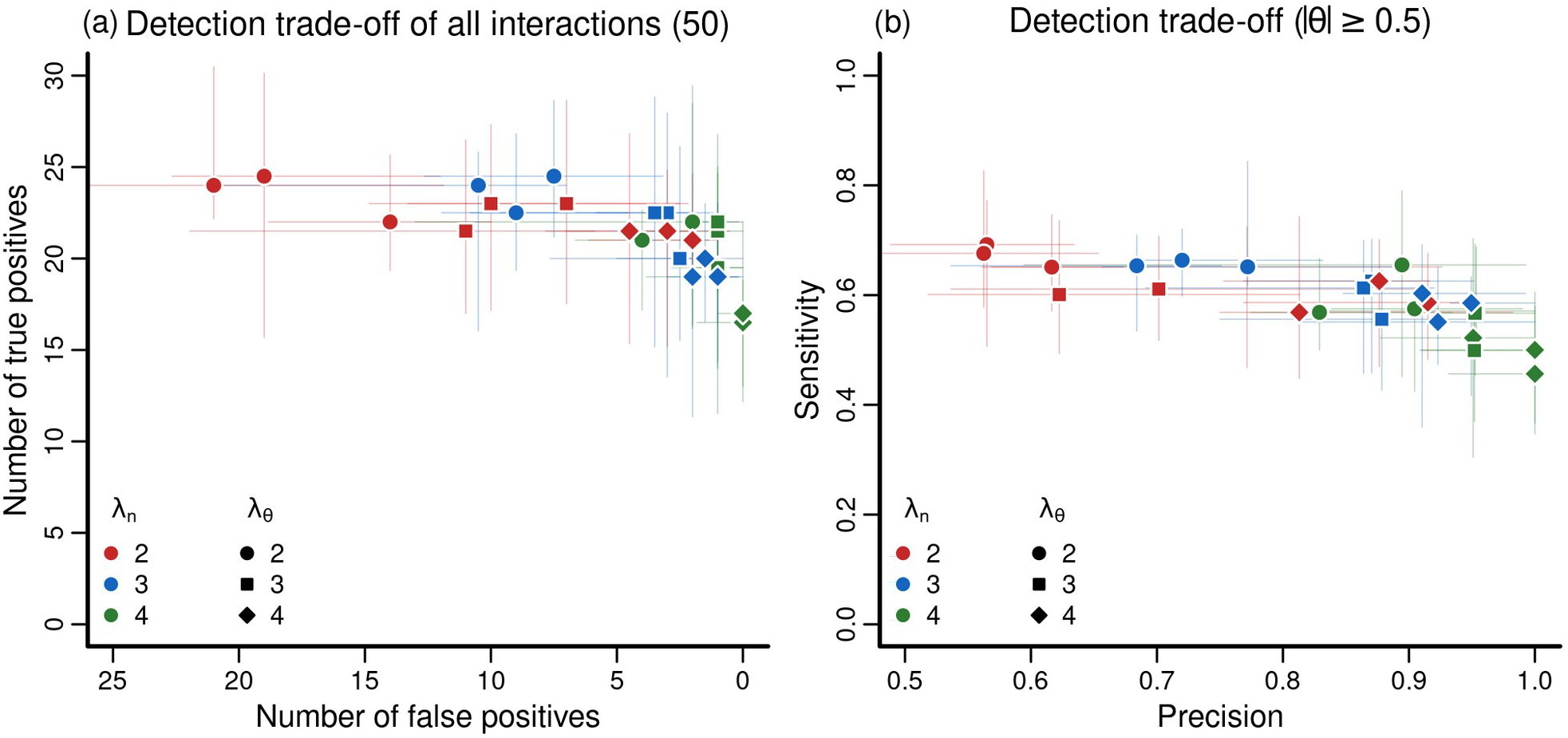
Accuracy of the SCEM model with two missing predictors (estimated as latent variables), in detecting true species interactions (y axis) while discarding spurious interactions (x axis) (a), and the precision-sensitivity trade-off in respect to interaction detection (b), for simulated communities with 64 species generated with 50 true species interactions, under different hyperparameterizations (colors and symbols). Colors and symbols depict λ_n_ and λ_θ_ respectively; the third varying hyperparameter (λ_β_) was not categorized because it had a smaller influence. Each dot-and-whiskers corresponds to one set of hyperparameter values and depicts the median and the 95% quantile range of 8 different models estimated and selected with that hyperparameterization. In (b), for sensitivity, we only counted undetected interactions whose absolute true value was higher than 0.5 as false negatives, thereby giving focus to the most serious false negatives (i.e. strong interactions that were not detected). For all calculations, we focused on the detection of interactions regardless of their direction, since the estimation accuracy of direction was analysed separately (Fig. S4). The total number of interactions is, therefore, 2016 (half of the θ matrix excluding diagonal).

In terms of the common classification model accuracy metrics for the recovered interactions, the model stands out for its very high specificity and accuracy in all tested hyperparameterizations, which comes at the cost of an intermediate sensitivity (Fig. 4, Fig. S2). The very high values of accuracy and specificity (always greater than 0.98) result from the very high numbers of true negatives – on average, around 1960 in the cases herein presented.

**Figure 4:**
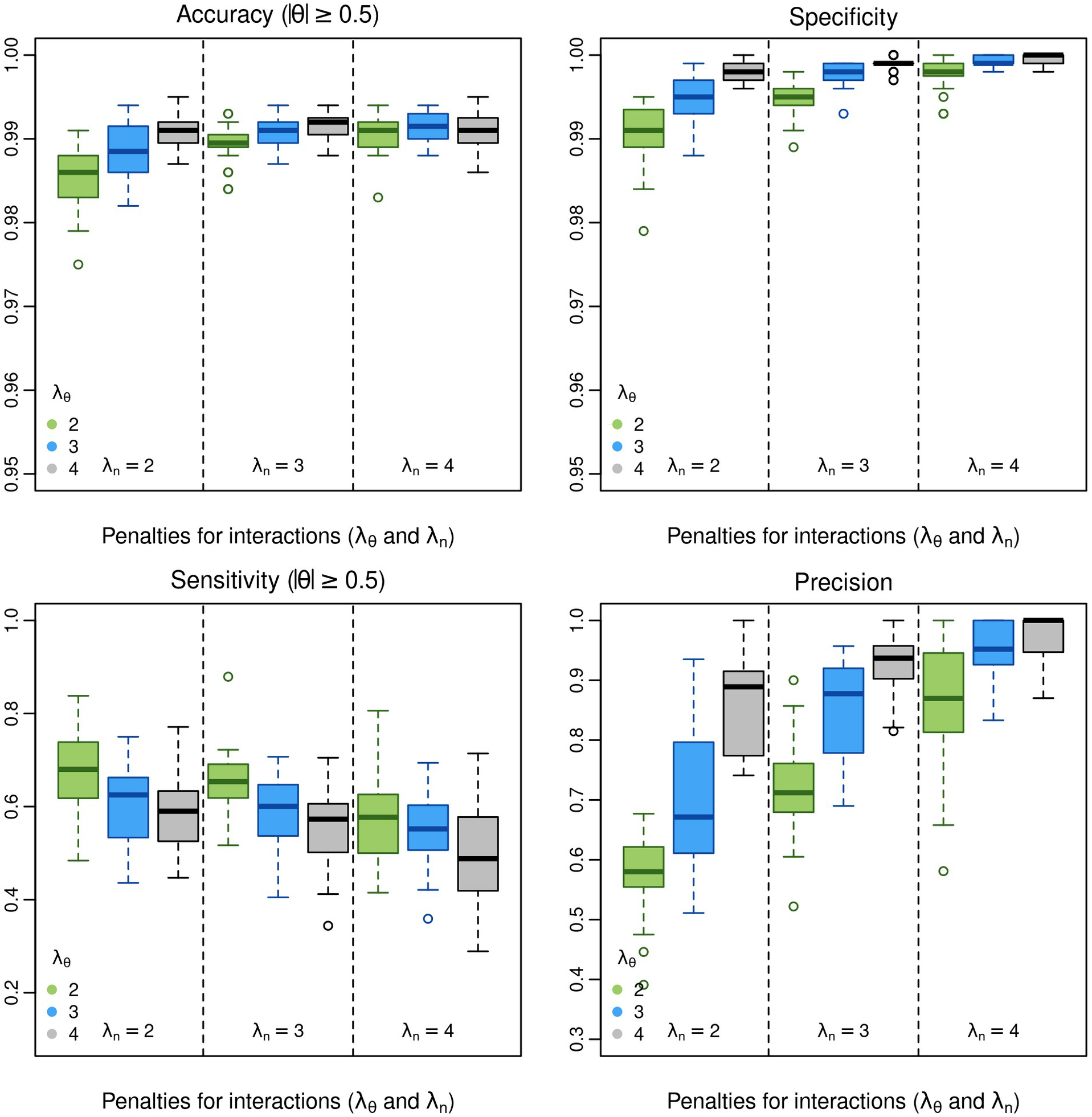
Model quality assessment (accuracy, specificity, sensitivity and precision) of the SCEM model with two missing predictors (estimated as latent variables), in respect to the identified species interactions, for simulated communities with 64 species generated with 50 true species interactions, under different hyperparameterizations (x axis). The x axis represents a combination of the penalty applied to the interaction coefficients λ_θ_ and the penalty of the number of interactions in the model λ_n_. For accuracy and sensitivity, we only counted undetected interactions whose absolute true value was higher than 0.5 as false negatives, thereby giving focus to the most serious false negatives (i.e. strong interactions that were not detected). For all calculations, we focused on the detection of interactions regardless of their direction, since the estimation accuracy of direction was analysed separately (Fig. S4).

Regarding the interaction strengths and signals of the recovered interactions, there was a very good match between the estimated interaction coefficients (θ) and their true values (Fig. S3a), with an R^2^ of about 95% between the true and estimated values, in all tested hyperparameterizations. Results were also very good for the environmental coefficients (Fig. S3b), although in this case, there was a marginal decreasing trend of the R^2^ as the penalty of environmental coefficients (λ_β_) was increased, reflecting the expected increasing bias due to heavier regularization. Likewise, the percent of true interactions that were recovered in the wrong direction (species A affects species B vs. B affects A) was lower than chance (Fig. S4), varying roughly from 20% to 40% of the detected interactions, and was apparently unaffected by the hyperparameterization.

Results for sample communities with 128 species and 100 interactions (Fig. 5) suggest that the model accuracy remains unchanged as compared to the 64-species communities, despite the major increase in model complexity, which has four times more possible interactions available to be selected (8128). The branched chains of indirect effects did not affect much the accuracy of interaction detection, in particular: the true positives were detected irrespectively of whether they belonged to long chains or not; and there was apparently no tendency for the false positives to shortcut indirect effects as one could expect, i.e. in cases where the true links are A-B-C, the model did not falsely select A-C (Fig. 5, left panels). In terms of the strength and signal of the identified interactions, there was also no change in accuracy, with R^2^ between the true and estimated values of around 0.97, and no apparent bias (Fig. 5, right panels). Interactions that were recovered in the opposite direction (relative to the truth), were equally well estimated otherwise (Fig. 5, blue circles). Estimated latent variables closely matched the values of the true predictors that were dropped from model fitting with no apparent bias, particularly in the 128-species case (Fig. S5).

**Figure 5:**
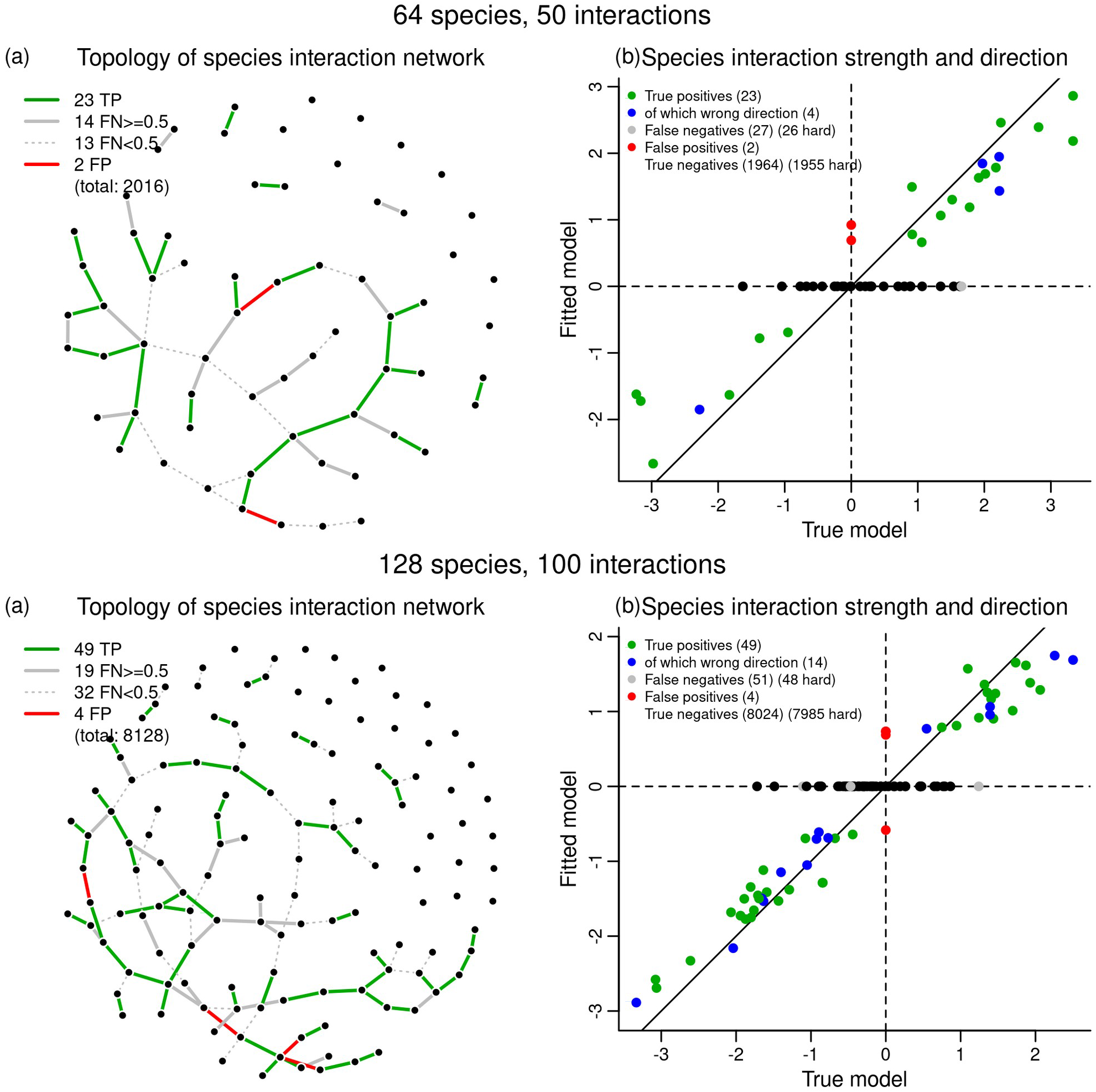
True versus fitted (a) species interaction network topology and (b) species interaction strength and direction, for two fitted models with 64 species-50 interactions and 128 species-100 interactions, with two missing predictors (estimated as latent variables). In (a), false negatives (gray lines) are split in weak interactions (dotted lines, absolute coefficient < 0.5) and strong interactions (solid lines, absolute coefficient >= 0.5). In (b), black circles depict the species interactions that were hard-thresholded before the network selection stage, whereas gray circles those that entered the selection stage but were not selected. TP: true positives, FN: false negatives, FP: false positives. Quality metrics (64 species): Accuracy = 0.99; Sensitivity = 0.46 (0.62, |θ| >= 0.5); Specificity ~ 1.00; Precision = 0.92; R^2^ of the true positives = 0.95. Quality metrics (128 species): Accuracy = 0.99; Sensitivity = 0.49 (0.72, |θ| >= 0.5); Specificity ~ 1.00; Precision = 0.93; R2 of the true positives = 0.97. Hyperparameters: λ_β_ = 5, λ_θ_ = 3, λ_n_ = 3, θ_0_ = 0.5, λ_υ_ = 1 (both models).

Altogether, results show that the model can correctly tell apart the effects of species interactions from those of shared responses to environment using as sole input plain presence/absence data where both effects are intermingled, under the scenario of missing predictors inducing confounding co-occurrence patterns. Furthermore, the model was able to recover very well the topology of the true interaction network after selection even in the most complex networks, which means that the network selection algorithm was capable of correctly excluding the great majority of otherwise false positives. This is noteworthy, given that in sparse networks as we tested, there are usually thousands of potential false positives. It is also interesting to note that the estimates for the true interactions under a full network model (i.e. without selection) closely matched the true values (Fig. S6), which shows that the model may produce accurate estimates even when obviously overfitted. This legitimates the idea of pruning out interactions whose preliminary estimates (i.e. under the full model) are weak, before conducting network selection, as a strategy for reducing model complexity.

### Limitations and future directions

Our framework based on Simultaneous Community Equations Models (SCEM) can contribute to enhance our understanding of community assembly mechanisms, by explicitly incorporating the effects of species interactions on community composition. Although performance comparisons across methods were beyond the scope of our paper, SCEM likely provides more accurate information than other methods, because the estimation of each interaction is conditional on all other possible interactions, and on the confounding effects of unmeasured drivers, neither of which are controlled through methods based for instance on association matrices (Blanchet *et al*. 2020). Moreover, by treating interactions explicitly, it is closer to a process-based model than models based on association patterns, where the interaction process itself is implicit (Dormann *et al*. 2012), hence, a source of misinterpretations. Like any correlative method, however, SCEM cannot be used to prove the actual functional significance of the interactions detected (Dormann *et al*. 2012; Ovaskainen *et al*. 2017a). Nonetheless, it provides a robust quantitative basis to identify such functionally important interactions, which should then be subjected to more detailed experimental scrutiny.

SCEM can be easily expanded to accommodate other types of community information, which have already been considered in community models, and are important to gain a more complete and precise understanding of community assembly mechanisms (Kissling *et al*. 2012; D’Amen *et al*. 2017). A straightforward expansion is to model abundance data rather than just presence/absence, which can be made simply by altering the likelihood function and allowing other link functions. Also, the model can be readily expanded to incorporate the effects of traits on the way species interact with the environment (Ovaskainen *et al.* 2017a), and account for multilevel sampling designs and spatial random effects (Ovaskainen *et al*. 2016). Also interesting would be to incorporate the effects of traits (or functional groups/guilds) on species interactions, but this is more challenging conceptually and technically (e.g., Kissling *et al*. 2012; Ovaskainen *et al*. 2017b). However, the flexibility of the model formulation and estimation mechanism, which is free from assumptions, approximations and simplifications, constitutes an advantage for implementing any of these extensions, by not requiring core changes in the penalized maximum likelihood optimization algorithm.

In summary, the SCEM framework offers a new way to incorporate species interactions in community models, while controlling for the confounding effects of unmeasured drivers and of all other possible interactions. By doing so, and by coupling that with the selection of network topology, it holds great potential to reveal biotic interaction networks from co-occurrence data, overcoming the long standing conundrum of mistaking shared niches for interactions (Ovaskainen *et al*. 2016; Ovaskainen *et al*. 2017a; Blanchet *et al*. 2020), something that was hampered by multiple shortcomings associated with other methodological approaches. Despite the limitations highlighted above, we have shown the usability of the model with a real use case and believe that the current advances already provide a valuable tool to enable ecologists to better tackle these more obscure processes of community assembly. At the same time, we hope that the limitations of the current implementation actually stimulate future developments upon our approach.

## Acknowledgements

We thank Tiago Marques and César Capinha for their comments on a first draft of the manuscript. MP was supported by national funds through FCT – Fundação para a Ciência e a Tecnologia, I.P. in the scope of Norma Transitória – DL57/2016/CP1440/CT0017, and PB by EDP Biodiversity Chair.

### Box 1: Why Simultaneous Equations Modelling can improve the inference of biotic interactions from community data?

Most community modelling methods that incorporate the effects of biotic interactions suffer from a structural limitation: they seek the signal of species interactions using symmetric species associations, which measure the probability of co-occurrence without accounting for the possible interactions with other species and for other processes that cause species associations (Blanchet *et al.* 2020). While such approach is practical from a statistical point of view, because the association matrix may be calculated from the responses of species to a small set of latent variables, it is not fully correct to infer biotic interactions.

For instance, consider the basic interaction whereby the shade of forest trees affect herb species growing underneath. Shade can be regarded as an unmeasured micro-environmental covariate (an “interaction currency”, Kissling *et al.* 2012; Warton *et al.* 2015), and modelled using a latent variable. The interactions might be inferred from the responses of species to this latent variable: both herbs and trees would “respond” to this latent variable, hence become associated in a residual correlation matrix due to having similar (or opposite) responses. However, the shade, is caused by the trees only, and is available to the understory herbs only, but the latent variable approach does not account for this asymmetry – response to a latent variable versus effect on the latent variable (Kissling *et al.* 2012). Therefore, it confounds the effects of interactions with the effects of shared responses to unmeasured environmental drivers. More importantly, because all herbs respond to this variable “shade”, they would become associated with each other as well, and the same from the trees. For example, all shade-loving herbs would be positively correlated (as well as the trees), and there would be no way to distinguish this pattern, reflecting false positive interactions, from the association of herbs with trees (reflecting true interactions) because such model does not control explicitly for the effects of other interactions. Although not based on latent variables, the same rationale applies to a more recent approach whereby the association matrix is parameterized directly without resourcing to latent variables (Pichler & Hartig 2020), as species association patterns would still reflect the associations that lead to the same inference of false positive interactions.

Simultaneous Equations Models allow to formulate the same problem differently: interactions may now be regarded as direct effects of each species on each of the others. Each species pair has, therefore, two possible interactions. Because all interactions are explicitly incorporated in the model as conventional model terms, the estimation of one particular interaction accounts for the effects of all the others, just like the estimation of a linear model coefficient accounts for the others. If we add the effects of latent variables to the system of simultaneous equations, the model will further control for the effects of other confounding processes also causing co-occurrence. This conceptual shift, therefore, offers a solution to a) discriminate between shared environmental responses and interactions (Blanchet *et al.* 2020). because the effects of species interactions are separated by construction from those of shared responses to unmeasured drivers; b) control for the effects of all other interactions when estimating an interaction, because all interactions are estimated simultaneously, therefore, the inference of an individual interaction accounts for all the others; c) estimate asymmetric interactions because interactions are modelled as a directed graph, hence the direction (asymmetry) of interactions is explicitly incorporated; d) recover interaction networks even with cycles, because the network is not solved as in the Markov and Bayesian network approaches (Harris 2016; Staniczenko *et al.* 2017), but estimated as if in an equilibrium state. An additional advantage of having explicit terms for species interactions is that it enables conducting variable selection on interactions and/or augmenting the model with previous ecological knowledge by excluding particular interactions *a priori*, which is not possible with association based methods.

## References

Abrego, N., Roslin, T., Huotari, T., Tack, A.J.M., Lindahl, B.D., Tikhonov, G. et al. (2020). Accounting for environmental variation in co-occurrence modeling reveals the importance of positive interactions in root-associated fungal communities. Mol. Ecol., 29, 2736–2746.

Baltagi, B.H. (2008). Simultaneous Equations Model. In: Econometrics. Springer-Verlag, Berlin, Heidelberg, pp. 257–303.

Bell, D.M. & Schlaepfer, D.R. (2016). On the dangers of model complexity without ecological justification in species distribution modeling. Ecol. Modell., 330, 50–59.

Blanchet, F.G., Cazelles, K. & Gravel, D. (2020). Co-occurrence is not evidence of ecological interactions. Ecol. Lett., 23, 1050–1063.

Calcagno, V. (2020). glmulti: Model Selection and Multimodel Inference Made Easy. R package version 1.0.8. Available at: https://cran.r-project.org/package=glmulti

Caradima, B., Schuwirth, N. & Reichert, P. (2019). From individual to joint species distribution models: A comparison of model complexity and predictive performance. J. Biogeogr., 46, 2260–2274.

Cazelles, K., Araújo, M.B., Mouquet, N. & Gravel, D. (2016). A theory for species co-occurrence in interaction networks. Theor. Ecol., 9, 39–48.

D’Amen, M., Rahbek, C., Zimmermann, N.E. & Guisan, A. (2017). Spatial predictions at the community level: from current approaches to future frameworks. Biol. Rev., 92, 169–187.

D’Amen, M., Mod, H.K., Gotelli, N.J. & Guisan, A. (2018). Disentangling biotic interactions, environmental filters, and dispersal limitation as drivers of species co-occurrence. Ecography, 41, 1233–1244.

Dormann, C.F., Schymanski, S.J., Cabral, J., Chuine, I., Graham, C., Hartig, F., et al. (2012). Correlation and process in species distribution models: bridging a dichotomy. J. Biogeogr., 39, 2119–2131.

Dormann, C.F., Bobrowski, M., Dehling, D.M., Harris, D.J., Hartig, F., Lischke, H., et al. (2018). Biotic interactions in species distribution modelling: ten questions to guide interpretation and avoid false conclusions. Glob. Ecol. Biogeogr., 27, 1004–1016.

Gotelli, N.J. & Ulrich, W. (2010). The empirical Bayes approach as a tool to identify non-random species associations. Oecologia, 162, 463–477.

Götzenberger, L., de Bello, F., Bråthen, K.A., Davison, J., Dubuis, A., Guisan, A., et al. (2012). Ecological assembly rules in plant communities-approaches, patterns and prospects. Biol. Rev. Camb. Philos. Soc., 87, 111–127.

Harris, D.J. (2016). Inferring species interactions from co-occurrence data with Markov networks. Ecology, 97, 3308–3314.

Heinze, G. & Schemper, M. (2002). A solution to the problem of separation in logistic regression. Stat. Med., 21, 2409–2419.

Hoerl, A.E. & Kennard, R.W. (1970). Ridge Regression: Biased Estimation for Nonorthogonal Problems. Technometrics, 12, 55–67.

Hui, F.K.C., Taskinen, S., Pledger, S., Foster, S.D. & Warton, D.I. (2015). Model-based approaches to unconstrained ordination. Methods Ecol. Evol., 6, 399–411.

Hutchinson, R.A., Valente, J.J., Emerson, S.C., Betts, M.G. & Dietterich, T.G. (2015). Penalized likelihood methods improve parameter estimates in occupancy models. Methods Ecol. Evol., 6, 949–959.

Kissling, W.D., Dormann, C.F., Groeneveld, J., Hickler, T., Kühn, I., Mcinerny, G.J., et al. (2012). Towards novel approaches to modelling biotic interactions in multispecies assemblages at large spatial extents. J. Biogeogr., 39, 2163–2178.

Levine, J.M., Bascompte, J., Adler, P.B. & Allesina, S. (2017). Beyond pairwise mechanisms of species coexistence in complex communities. Nature, 546, 56–64.

Morales-Castilla, I., Matias, M.G., Gravel, D. & Araújo, M.B. (2015). Inferring biotic interactions from proxies. Trends Ecol. Evol., 30, 347–356.

Nieto-Lugilde, D., Maguire, K.C., Blois, J.L., Williams, J.W. & Fitzpatrick, M.C. (2018). Multiresponse algorithms for community-level modelling: Review of theory, applications, and comparison to species distribution models. Methods Ecol. Evol., 9, 834–848.

Niku, J., Hui, F.K.C., Taskinen, S. & Warton, D.I. (2019). gllvm: Fast analysis of multivariate abundance data with generalized linear latent variable models in R. Methods Ecol. Evol., 10, 2173–2182.

Niku, J., Warton, D.I., Hui, F.K.C. & Taskinen, S. (2017). Generalized linear latent variable models for multivariate count and biomass data in ecology. J. Agric. Biol. Environ. Stat., 22, 498–522.

Norberg, A., Abrego, N., Blanchet, F.G., Adler, F.R., Anderson, B.J., Anttila, J., et al. (2019). A comprehensive evaluation of predictive performance of 33 species distribution models at species and community levels. Ecol. Monogr., 89, e01370.

Ovaskainen, O. & Soininnen, J. (2011). Making more out of sparse data: hierarchical modeling of species communities. Ecology, 92, 289–295.

Ovaskainen, O., Abrego, N., Halme, P. & Dunson, D. (2016). Using latent variable models to identify large networks of species-to-species associations at different spatial scales. Methods Ecol. Evol., 7, 549–555.

Ovaskainen, O., Tikhonov, G., Norberg, A., Guillaume Blanchet, F., Duan, L., Dunson, D., et al. (2017a). How to make more out of community data? A conceptual framework and its implementation as models and software. Ecol. Lett., 20, 561–576.

Ovaskainen, O., Tikhonov, G., Dunson, D., Grøtan, V., Engen, S., Sæther, B.E., et al. (2017b). How are species interactions structured in species-rich communities? A new method for analysing time-series data. Proc. R. Soc. B Biol. Sci., 284.

Pichler, M. & Hartig, F. (2020). A new method for faster and more accurate inference of species associations from novel community data. arXiv preprint arXiv:2003.05331v4.

Pollock, L.J., Tingley, R., Morris, W.K., Golding, N., O’Hara, R.B., Parris, K.M., et al. (2014). Understanding co-occurrence by modelling species simultaneously with a Joint Species Distribution Model (JSDM). Methods Ecol. Evol., 5, 397–406.

Popovic, G.C., Warton, D.I., Thomson, F.J., Hui, F.K.C. & Moles, A.T. (2019). Untangling direct species associations from indirect mediator species effects with graphical models. Methods Ecol. Evol., 10, 1571–1583.

Porto, M. & Beja, P. (2020). eicm: Explicit Interaction Community Models. Available at: https://cran.r-project.org/package=eicm

Rothenberg, T.J. (1990). Simultaneous Equations Models. In: Econometrics (eds Eatwell, J., Milgate, M. & Newman, P.). Palgrave Macmillan UK, London, pp. 229–237.

Sander, E.L., Wootton, J.T. & Allesina, S. (2017). Ecological Network Inference from Long-Term Presence-Absence Data. Sci. Rep., 7, 7154.

Soliveres, S., Maestre, F.T., Ulrich, W., Manning, P., Boch, S., Bowker, M.A., et al. (2015). Intransitive competition is widespread in plant communities and maintains their species richness. Ecol. Lett., 18, 790–798.

Staniczenko, P.P.A., Sivasubramaniam, P., Suttle, K.B. & Pearson, R.G. (2017). Linking macroecology and community ecology: refining predictions of species distributions using biotic interaction networks. Ecol. Lett., 20, 693–707.

Tikhonov, G., Duan, L., Abrego, N., Newell, G., White, M., Dunson, D. et al. (2020a). Computationally efficient joint species distribution modeling of big spatial data. Ecology, 101, e02929.

Tikhonov, G., Opedal, Ø.H., Abrego, N., Lehikoinen, A., de Jonge, M.M.J., Oksanen, J., et al. (2020b). Joint species distribution modelling with the R-package Hmsc. Methods Ecol. Evol., 11, 442–447.

Tobler, M.W., Kéry, M., Hui, F.K.C., Guillera-Arroita, G., Knaus, P. & Sattler, T. (2019). Joint species distribution models with species correlations and imperfect detection. Ecology, 100, e02754.

Warton, D.I., Blanchet, F.G., O’Hara, R.B., Ovaskainen, O., Taskinen, S., Walker, S.C., et al. (2015). So Many Variables: Joint Modeling in Community Ecology. Trends Ecol. Evol., 30, 766–779.

Wilkinson, D.P., Golding, N., Guillera-Arroita, G., Tingley, R. & McCarthy, M.A. (2019). A comparison of joint species distribution models for presence–absence data. Methods Ecol. Evol., 10, 198–211.

Wisz, M.S., Pottier, J., Kissling, W.D., Pellissier, L., Lenoir, J., Damgaard, C.F., et al. (2013). The role of biotic interactions in shaping distributions and realised assemblages of species: Implications for species distribution modelling. Biol. Rev., 88, 15–30.

Zurell, D., Pollock, L.J. & Thuiller, W. (2018). Do joint species distribution models reliably detect interspecific interactions from co-occurrence data in homogenous environments? Ecography, 41, 1812–1819.

