## Supporting Information for "Simultaneous equations modelling of communities with interacting species networks"

**This document contains four appendices and seven figures.**

### Appendix S1. Fitting workflow for Simultaneous Community Equations Models

The Linear Predictor of the Simultaneous Community Equations Models takes the form

$$LP=\begin{pmatrix} X_{11} & X_{12} & \cdots & X_{1k} & \upsilon_{11} & \upsilon_{12} & \cdots & \upsilon_{1u} \\ X_{21} & X_{22} & \cdots & X_{2k} & \upsilon_{21} & \upsilon_{22} & \cdots & \upsilon_{2u} \\ X_{31} & X_{32} & \cdots & X_{3k} & \upsilon_{31} & \upsilon_{32} & \cdots & \upsilon_{3u} \\ \vdots & \vdots & \ddots & \vdots & \vdots & \vdots & \ddots & \vdots \\ X_{m1} & X_{m2} & \cdots & X_{mk} & \upsilon_{m1} & \upsilon_{m2} & \cdots & \upsilon_{mu} \end{pmatrix} \times \begin{pmatrix} \beta_{11} & \beta_{12} & \cdots & \beta_{1n} \\ \beta_{21} & \beta_{22} & \cdots & \beta_{2n} \\ \vdots & \vdots & \ddots & \vdots \\ \beta_{(k+u)1} & \beta_{(k+u)2} & \cdots & \beta_{(k+u)n} \end{pmatrix} + \begin{pmatrix} Y_{11} & Y_{12} & \cdots & Y_{1n} \\ Y_{21} & Y_{22} & \cdots & Y_{2n} \\ Y_{31} & Y_{32} & \cdots & Y_{3n} \\ \vdots & \vdots & \ddots & \vdots \\ Y_{m1} & Y_{m2} & \cdots & Y_{mn} \end{pmatrix} \times \begin{pmatrix} 0 & \theta_{12} & \cdots & \theta_{1n} \\ \theta_{21} & 0 & \cdots & \theta_{2n} \\ \vdots & \vdots & \ddots & \vdots \\ \theta_{n1} & \theta_{n2} & \cdots & 0 \end{pmatrix}$$

$= [X\ \upsilon] \times \beta + Y \times \theta$

$= X \times \beta^X + \upsilon \times \beta^\upsilon + Y \times \theta$

where n is the number of species, m is the number of sampling sites, k is the number of measured environmental predictors and u is the number of unknown environmental predictors. X are the measured environmental variables at each site, υ are the unmeasured predictor values at each site, β are the effects of each environmental predictor (measured: $\beta^X$ and unmeasured: $\beta^\upsilon$) on each species, Y the presence/absence (or abundance) of each species at each site, and θ are the effects of each species on each of the other species. X and Y are the known data (note that the X submatrix may be missing), whereas υ, β and θ are unknowns to be estimated.

The above formulation is rotational invariant in respect to υ and $\beta^\upsilon$ (the rows of β corresponding to the columns υ) when there are latent variables to be estimated (u > 0) because there are infinite combinations of υ and $\beta^\upsilon$ yielding the same value of the LP, and hence, rendering the model non-identifiable.

*Proof of rotational invariance of $\upsilon \times \beta^\upsilon$*

As a proof, we want to show that rotating υ and rotating the corresponding $\beta^\upsilon$ with the same rotation matrix R yields the same LP. Note that, to rotate, we have to transpose $\beta^\upsilon$ so that the latent variables are along the columns, and, after rotation, back-transpose to put back the latents along the rows. So, we want to show that

$\upsilon \times \beta^\upsilon = (\upsilon \times R) \times (\beta^{\upsilon T} \times R)^T$ which implies

$\upsilon \times \beta^\upsilon = \upsilon \times R \times R^T \times \beta^\upsilon$

Since $R^T = R^{-1}$ because R is a rotation matrix, this is equivalent to

$\upsilon \times \beta^\upsilon = \upsilon \times R \times R^{-1} \times \beta^\upsilon$ and hence,

$\upsilon \times \beta^\upsilon = \upsilon \times \beta^\upsilon$

Since $[X\ \upsilon] \times \beta = X \times \beta^X + \upsilon \times \beta^\upsilon$, it follows that the $[X\ \upsilon] \times \beta$ term of the LP remains constant irrespectively of the rotation applied to both υ and $\beta^\upsilon$.

#### Model fitting and selection

Due to the rotational invariance of $\upsilon \times \beta^\upsilon$, it is not feasible to estimate all model parameters at once without applying a constraint (Warton *et al.* 2015). In particular, the estimation of interactions (θ) requires the rotational invariance to be fixed. To achieve that, the values of the latent variables at each site υ – but not the respective species responses – can be estimated and fixed prior to the estimation of the species interactions, thereby cancelling the rotational invariance. For the purposes of estimating species interactions, it is irrelevant which rotation is chosen, because they all yield the same LP, hence do not affect the estimation of the interaction terms.

However, as detailed in the main text, the estimation of the full model (all non-diagonal elements of θ) is barely informative from an ecological viewpoint because of the large amount of spurious species interactions that are estimated in θ. To minimize this problem, we developed a selection algorithm upon the θ parameters (which make up the topology of the interaction network), which is described in the main text. However, the number of parameters in the θ matrix increases quadratically in respect to the number of species, which may pose serious difficulties when conducting the network topology selection. It is, therefore, advisable to reduce network complexity before entering the selection stage.

To reduce the complexity of the network selection, and thus increase the chance of finding the global most parsimonious interaction network, we propose that the selection is conducted in a reduced model, i.e. using a subnetwork as the "full" model. There are three major options to do this, the first (a) is conducted by default in the fitting function, but the user may additionally opt for the other two, which are also implemented:

a) weak interactions may be discarded *via* hard thresholding (with a threshold $|\theta| < \theta_0$) upon the initial, full network estimates. This choice is generally reasonable and very effective even for small values of $\theta_0$ (e.g. $\theta_0 = 0.5$), because a great part of full model interaction estimates are close to zero, owing to LASSO regularization.

b) impossible or very unlikely interactions may be excluded *a priori* (Wisz *et al*. 2013; Morales-Castilla *et al*. 2015), for example, based on species traits or other type of ecological knowledge, thus augmenting model estimation with ecological knowledge. This is usually straightforward at least for specifying direction, as it is generally easy, given a pair of species (or guilds/functional groups), to ascertain in which direction the eventual interaction is stronger. For example, in a plant dataset, we may assume that the influence that trees may have on herbs is much stronger than the opposite (e.g. through shading), and so we may specify a priori that herbs do not affect trees (but the reverse), thereby much reducing model complexity. The 'eicm' package fitting function has simple ways for specifying such trait-based or species-based exclusions.

c) interactions departing from rare species may be excluded (Le Roux *et al*. 2014) because, even though rare species may have important impacts on others, their reduced number of occurrences will limit the statistical detectability of their effects. This exclusion is also easily conducted with the fitting function.

Under the setting described above, the proposed complete fitting workflow, that is implemented in the main fitting function of the 'eicm' package, is as follows:

Step 1. Estimate the latent variable matrix υ (i.e. the values of missing predictors for each site), assuming no species interactions. Fix these values for the next steps;

Step 2. Estimate species interaction coefficients θ under a full network model (all species interactions allowed except those excluded *a priori* as per the user-defined criteria) with the latent variable values υ fixed (as in Step 1);

Step 3. Discard from the full model (i.e. fix at 0) all species interactions with estimated absolute value (in Step 2) lower than a threshold $\theta_0$ – this constitutes the reduced model;

Step 4. With the latent variable values $\upsilon$ fixed (as in Step 1) – but not their respective species coefficients –, conduct variable selection on the species interactions using the reduced network model (Step 3) as the full model. Note that the interaction coefficients that were estimated in Step 2 are discarded, and obviously re-estimated for each candidate species interaction network topology.

#### Setting model hyperparameters

There are five hyperparameters, plus the number of latent variables to estimate, that must be set along the fitting workflow. These control the trade-offs between tractability, errors of Type I and II, and bias during the model fitting process. The hyperparameters are:

$\lambda_\upsilon$: the penalty applied to latent variables and species environmental coefficients in Step 1;

$\lambda_\beta$: the penalty applied to species environmental coefficients in all the remaining steps;

$\lambda_\theta$: the penalty applied to species interaction coefficients in all the remaining steps;

$\theta_0$: the cutoff threshold used to discard weak species interactions from the full model in Step 3;

$\lambda_n$: the weight applied to the penalty of the number of interaction terms included during the variable selection step (Step 4).

Two important points of concern to the end users are how to set these model hyperparameters and how to define an adequate number of latent variables to estimate. Both problems are common to other community models (e.g. Ovaskainen *et al*. 2016; Pichler & Hartig 2020). Regularization penalties are usually tuned with cross-validation, by maximizing the likelihood in test data (Hutchinson *et al*. 2015; Sander *et al*. 2017; Pichler & Hartig 2020), or, more generally, any set of hyperparameters can be optimized by cross-validation with heuristics (Vignali *et al*. 2020). However, cross-validation is not applicable for the problem that we focus in this paper – maximize the accuracy of biotic interaction detection – because there is no empirical data on species interactions on which to test models against: the only “criterion” that can be evaluated in this respect is the ecological meaningfulness of detected interactions.

For the inference of biotic interactions, the accuracy screening of different penalties we have conducted with virtual communities, hence, provides important guidance (Fig. 3-4 in the main text, Fig. S2). This extensive trialling showed that $\lambda_\theta$ and $\lambda_n$ are the most influent on the trade-off between Type I and Type II errors, whereas $\theta_0$ is fundamental not only for controlling the algorithm running time but also its efficiency, as the more spurious interactions that are discarded from the start, the more likely the algorithm is to find the “true” interactions, at the expense of discarding some “true” weak interactions. The other hyperparameters have shown to be less important and may be set to their defaults, which are small values whose purpose is to avoid parameter divergence in otherwise flat likelihood surfaces.

Yet, results with real data (Appendix S2) suggest that the direct application of the simulation results to real use cases was not a perfect option, as the interaction penalties had to be decreased to allow their detection, which is probably because in real communities the interaction signals are generally weaker. We suggest that our simulation results are used as a starting point, but the user must critically evaluate results from an ecological viewpoint and revise his decisions appropriately. This may involve not only adjusting model hyperparameters, but also a better refinement of the a priori restrictions applied to the interaction network (Morales-Castilla *et al*. 2015).

As to the number of latent variables to include, this is also a problem that is shared with other community models. There, it is either itself subjected to regularization and truncation during model estimation, using a specific prior in a Bayesian framework (Bhattacharya & Dunson 2011; Warton *et al*. 2015; Ovaskainen *et al*. 2016), or no method is available to tune its value (Niku *et al*. 2017), as is our case also. An option to have an initial value for this setting would be to run a (partial) Principal Component Analysis (Oksanen *et al*. 2019) on the occurrence matrix and use the cumulative proportion of variance as a criterion for deciding how many latent variables would be enough to account for residual species co-variation. In any case, if the selected model includes an obvious excess of detected interactions, this may be indicative that not all the co-variation attributable to missing predictors was accounted for, and more latent variables may be needed.

### Appendix S2. Case study

#### Study area

We illustrate the use of the SCEM model with a real dataset of 300 plant species whose presence or absence was recorded in 109 sampling sites. The sampling sites were located in natural forest patches dispersed within about 27,000 ha located in north-eastern Portugal, in a region with a Mediterranean climate. There are important climatic and microclimatic gradients due to the ruggedness of the landscape, which apparently cause marked differences in the dominant species, from herbs to trees. Hence, forests are dominated by combinations of *Quercus rotundifolia* (holm oak), *Quercus suber* (cork oak), *Juniperus oxycedrus*, and *Olea europaea* var. *sylvestris* (wild olive), with a varied understory including *Pistacia terebinthus* (turpentine tree), *Lavandula pedunculata*, *Erica arborea* (tree heath) and species of *Cytisus* (brooms) and *Cistus* (rockroses), among many other shrubs. The presence of all vascular plant species was recorded in three 2-meter radius circles per site, and aggregated at the site level. Species whose presence was probably not of natural origin were excluded (5 species), as well as records that could not be identified with certainty to species level (11 taxa). Data is available from the Dryad Digital Repository: http://dx.doi.org/10.5061/dryad.XXXXX.

#### Methods

As explanatory variables, we compiled three spatial climatic variables and bioclimatic indices with a resolution of 111 m from Monteiro-Henriques *et al*. (2015) and three topographical variables computed from a Digital Elevation Model with a resolution of 30 meters:

| **Variable** | **Description** |
|---|---|
| Positive Precipitation | Sum of the monthly precipitation of the months with positive mean temperature |
| Mean temperature of the warmest month | Maximum monthly average temperature |
| Ombrothermic index of the warmest bimonth of the summer quarter | Sum of the monthly precipitation of the two warmest months divided by the sum of the monthly average temperature of the same months |
| Slope | Angle between the tangent plane at the center of the sampling plot and the horizontal plane, expressed as the tangent. |
| Aspect (cosine) | Cosine of the projected angle between the normal vector of the tangent plane, and the north direction. Quantifies the north-south gradient of exposure to sun. |
| Aspect (sine) | Sine of the projected angle between the normal vector of the tangent plane, and the north direction. Quantifies the east-west gradient. |

Topographical variables were used as surrogates for local ecological conditions related to insolation, and to soil water and nutrient availability (Liu *et al*. 2014), such that steeper sites are more strongly limited by the latter two factors. Values for climatic and topographic variables used in models were extracted for the centre coordinates of each plot. All the variables were centered and scaled to unit variance.

With the presence/absence matrix and the environmental variable matrix, we fitted and selected the SCEM model with six latent variables. To avoid an exceedingly complex interaction network model, we only included interactions caused by trees and shrubs on all the oher lifeforms and within themselves. Additionally, we excluded species interactions caused by species with 10 occurrences or less (but not the reverse). The SCEM model was fitted with the following hyperparameterization: $\lambda_\beta = 5$, $\lambda_\theta = 2$, $\lambda_n = 1.5$, $\theta_0 = 0.5$, $\lambda_\upsilon = 1$.

To evaluate the quality of the estimated latent variables at each site, we compared their values with the site scores from a partial Principal Component Analysis conducted on the occurrence matrix, i.e. a PCA on the residuals after removing the effects of the six environmental variables on species occurrences (Oksanen *et al*. 2019). Under this setting, the axes extracted from the pPCA depict the most important community gradients expunged from the effects of the measured environmental variables, being therefore comparable, in principle, to the latent variables estimated in the SCEM. However, because SCEM latent factors are not ordered in terms of their explained variation (as in PCA), in order to be compared with the PCA axes, the best matching pairs were sought by means of a Pearson correlation matrix.

### Results

The fitted SCEM model suggests that our local communities were strongly influenced by environmental filters, having a AUC, of the whole community, of 0.94, for a null (intercepts only) AUC of 0.83, but that community assembly appeared to be influenced also by biotic interactions. 28% of the species did not show a strong response to any of the six measured environmental predictors, having all the respective coefficients lower than 0.25 in absolute value. The latent variables captured an important component of the unexplained variation, lowering the number of nearly unresponsive species to only 2%. The relevance of the latent variables was also supported by their spatial patterns, since all the six showed important spatial structures (Fig. S1). These are probably depicting more local scale abiotic factors to which the community is responding, and that were not accounted for by the measured covariates, which have a coarser resolution. Examples of such factors include tree cover, past and present land use, soil properties that depend on the bedrock (chemical composition, water retention, etc.), among others that were not included in the model. There was a good correspondence between the estimated latent variables and the site scores of a partial Principal Component Analysis applied to the occurrence data (Fig. S7). Although the order of the latent variables recovered by the SCEM is arbitrary, the best match of each one of the six latent variables was found in the first six pPCA axes.

The biotic effects were supported by 11 interactions retained after the selection procedure from a universe of 7200 possible interactions (accounting for direction) in the full restricted interaction network (Fig. 2). These interaction terms indicate that the probability of a species to be present in the community is affected (positively or negatively) by the presence of another species, even after accounting for the effects of measured and unmeasured (latent) environmental drivers. For instance, the model identified a negative effect of cork oak (*Quercus suber*) on the tiny understorey fern *Anogramma leptophylla*, even after accounting for the opposite responses of both species to the latent variable #1 (Fig. 2). This means that their mutually exclusive co-distribution pattern could not be sufficiently explained by opposite responses to a latent gradient, requiring a negative species interaction to explain the pattern. The inverse also happens, for example, *Rubus ulmifolius* tends to respond in opposite direction to latent variable #1 compared with *Brachypodium sylvaticum*, but there is a positive effect of the former over the latter, on top of that inverse pattern. These results show that the responses to unmeasured predictors are not necessarily correlated with the recovered species interactions, which contrasts with the usual approach, where interactions are estimated from the responses to unmeasured predictors. Like in any modelling approach, however, interpretations need to be careful and require ancillary information on for instance species ecology and natural history, as statistical relations do not necessarily imply causality. For instance, the positive effect of *Asparagus acutifolius* on *Pistacia terebinthus* is difficult to interpret and may reflect a spurious result, due for instance to shared habitat conditions that did not emerge with the six latent factors.

### Appendix S3. Model testing with simulated data (detailed methods)

The critical question to validate the model is to what extent the proposed model and fitting workflow (Appendix S1) is able to recover the true species interaction network (topology, including direction and interaction strengths) from simple presence-absence community data, in the presence of unmeasured predictors that induce confounding co-occurrence patterns. For this, we conducted a battery of trials with simulated data under different conditions, to assess model accuracy in recovering the interaction network. In each trial, we:

1. Simulated community data (a presence-absence matrix) generated with a known, sparse, interaction network and known environmental responses;
2. fitted a model discarding all the environmental predictors used for community data generation, thereby creating confounding effects in the form of shared responses to “unmeasured predictors”;
3. compared the fitted model with the true model (the one used in data generation), focusing on the topologies of the true and estimated interaction networks, after variable selection, and coefficients.

To make the testing environment more similar to real use cases, in addition to discarding all the environmental predictors, we conducted all analyses in a worst-case setting, namely:

a) we ensured that the simulated presence-absence data matrix followed a species abundance curve akin of that of real communities – i.e. where rare species clearly dominate;

b) we estimated the models with no prior knowledge about the interaction network, thus assuming that all interactions were possible.

This worst-case setting aimed to render the test results more comparable to reality by mimicking the usual situation in ecological studies, where important environmental predictors are often unknown/unmeasured, most species are rare and nothing is known about how they interact.

#### Methods

##### *Accuracy screening for different hyperparameterizations*

We first conducted a screening of model accuracy trade-offs for different model hyperparameterizations, using 64-species virtual communities with 50 species interactions. We screened different combinations of: (i) the penalty applied to the environmental coefficients during estimation, $\lambda_\beta$; (ii) the penalty applied to species interaction coefficients during estimation, $\lambda_\theta$; (iii) the weighting factor applied to the number of interaction terms in the selection stage, $\lambda_n$. Each hyperparameterization was repeated 8 times with different true models (i.e. we did 8 independent trials for each hyperparameterization), so that a measure of variation of the accuracy metrics, unbiased by the particular true model, could be calculated. $\theta_0$ was set to 0.5 and $\lambda_\upsilon$ set to 1 in all trials. Each trial consisted in the following steps:

#### Data generation

In each trial, simulated data (a site × species presence-absence matrix) was generated by one random realization of a parameterized model (the true model) for 64 species in 1000 samples. The true model was itself randomly generated in each trial, and included two environmental predictors (randomly drawn from a gaussian distribution) to which species responded with random strengths (see below). The true model also included 50 species interactions with random strengths (see below), with the constraint of representing a Directed Acyclic Graph. This constraint is necessary when realizing the model (but not for model fitting) because the probability of occurrence of a given species depends directly on other species' presences, which cannot be solved in the presence of cycles. In the absence of a closed form expression for predicting species probabilities, to realize communities, we conducted a simple staged process, which is implemented in the prediction function of the 'eicm' package: it starts by realizing species occurrences of those species that only depend on environmental predictors. Then, iteratively, continues realizing remaining species occurrence vectors of those species that only depend on environmental predictors and on species whose occurrence vectors have already been realized, until no species remain.

Because the species frequency distribution of biological communities follows the well-known pattern of many rare species and few common species, for the sake of simulated data realism, the true model was not parameterized completely randomly, but in a way that ensured that the frequency distribution (of species in samples) of the realized communities followed a given Beta distribution as much as possible (Marquet *et al.* 2017), using an heuristic optimization. The true model parameters, hence, did not follow any distribution, but the corresponding frequency distribution of the realized community did. The target Beta distribution was defined with the parameters shape1=1.5 and shape2=3. This model generation procedure is implemented in the utility function 'generateEICM' of the R package, and further details are supplied in the respective section of the manual.

#### Model fitting (estimation and selection)

We fitted the SCEM model according to the proposed model fitting workflow (Appendix S1) and using the defined hyperparameters for each trial. We tested all 27 combinations of the hyperparameters $\lambda_\beta \in \{5, 6, 7\}$; $\lambda_\theta \in \{2, 3, 4\}$; $\lambda_n \in \{2, 3, 4\}$. We used as sole model input the site × species presence-absence matrix, since the two environmental predictors used for generating the simulated data were discarded. In all models, we included the estimation of two latent variables.

#### Model accuracy assessment

The results of the best model obtained from the network selection, in each trial, were compared with the respective true model, in particular the parameter values (environmental and species interaction coefficients) and the network topology. The selected interaction network topology was compared with the true one by computing the common metrics of classification model accuracy applied to the presence or absence of interaction in the true and fitted networks, regardless of the coefficient values and direction, whose accuracy was evaluated separately. As false negatives, used in the calculation of sensitivity and accuracy, we only counted undetected interactions whose absolute true value was higher than 0.5, as a means to focus on the more serious false negatives (strong interactions that were not detected). Within the true positives (correctly identified interactions), we separated those interactions that were estimated in the correct direction from those estimated in the wrong (opposite) direction, to assess the accuracy in identifying the true direction. The accuracy of the estimated values of the environmental and species interaction coefficients was evaluated by computing the $R^2$ of a linear model modelling the fitted coefficients as a function of the true coefficients. In the case of the environmental coefficients, the linear model included, for each estimated latent variable, both true predictors because of the rotational invariance of the latent variables that precludes a 1:1 match between latents and true predictors.

##### *Accuracy of the model for larger communities*

We then conducted the above procedure to simulated communities with 128 species-100 interactions, but due to the computational burden involved, we did not run a comprehensive screening as above, but conducted isolated trials. The hyperparameters were set according to the best screening results, with adjustments where necessary. We analysed in more detail the topology and estimates of sample models, by plotting the true vs. fitted interaction networks and parameter values.

### Appendix S4. Supplementary references


Bhattacharya, A. & Dunson, D.B. (2011). Sparse Bayesian infinite factor models. *Biometrika*, 98, 291–306.

Hutchinson, R.A., Valente, J.J., Emerson, S.C., Betts, M.G. & Dietterich, T.G. (2015). Penalized likelihood methods improve parameter estimates in occupancy models. *Methods Ecol. Evol.*, 6, 949–959.

Liu, J., Yunhong, T. & Slik, J.W.F. (2014). Topography related habitat associations of tree species traits, composition and diversity in a Chinese tropical forest. *For. Ecol. Manage*., 330, 75–81.

Marquet, P.A., Espinoza, G., Abades, S.R., Ganz, A. & Rebolledo, R. (2017). On the proportional abundance of species: Integrating population genetics and community ecology. *Sci. Rep*., 7, 16815.

Monteiro-Henriques, T., Martins, M.J., Cerdeira, J.O., Silva, P., Arsénio, P., Silva, A., et al. (2015). Bioclimatological mapping tackling uncertainty propagation: Application to mainland Portugal. *Int. J. Climatol*., 411, 400–411.

Morales-Castilla, I., Matias, M.G., Gravel, D. & Araújo, M.B. (2015). Inferring biotic interactions from proxies. *Trends Ecol. Evol.*, 30, 347–356.

Niku, J., Hui, F.K.C., Taskinen, S. & Warton, D.I. (2019). gllvm: Fast analysis of multivariate abundance data with generalized linear latent variable models in R. *Methods Ecol. Evol.*, 10, 2173–2182.

Oksanen, J., Blanchet, F.G., Friendly, M., Kindt, R., Legendre, P., McGlinn, D., et al. (2019). *vegan: Community Ecology package*. Available at: https://cran.r-project.org/package=vegan

Ovaskainen, O., Abrego, N., Halme, P. & Dunson, D. (2016). Using latent variable models to identify large networks of species-to-species associations at different spatial scales. *Methods Ecol. Evol.*, 7, 549–555.

Pichler, M. & Hartig, F. (2020). A new method for faster and more accurate inference of species associations from novel community data. arXiv preprint arXiv:2003.05331v4.

Le Roux, P.C., Pellissier, L., Wisz, M.S. & Luoto, M. (2014). Incorporating dominant species as proxies for biotic interactions strengthens plant community models. *J. Ecol*., 102, 767–775.

Sander, E.L., Wootton, J.T. & Allesina, S. (2017). Ecological Network Inference from Long-Term Presence-Absence Data. *Sci. Rep.*, 7, 7154.

Vignali, S., Barras, A.G., Arlettaz, R. & Braunisch, V. (2020). SDMtune: An R package to tune and evaluate species distribution models. *Ecol. Evol*., 10, 11488–11506.

Warton, D.I., Blanchet, F.G., O'Hara, R.B., Ovaskainen, O., Taskinen, S., Walker, S.C., *et al*. (2015). So Many Variables: Joint Modeling in Community Ecology. *Trends Ecol. Evol.*, 30, 766–779.

Wisz, M.S., Pottier, J., Kissling, W.D., Pellissier, L., Lenoir, J., Damgaard, C.F., *et al*. (2013). The role of biotic interactions in shaping distributions and realised assemblages of species: Implications for species distribution modelling. *Biol. Rev.*, 88, 15–30.

**Figure S1.** Estimated latent variables of an SCEM fitted to real data (graduated from green to red), for each sampling site, projected on their spatial location. The curves represent the predicted values of a Generalized Additive Model modelling each latent variable as a function of the coordinates (WGS84 / UTM zone 29N), smoothed with a spline interaction smoother with a basis dimension of 20. The respective $R^2$ is displayed. See main text and Appendix S2 for details on the real data analysis.

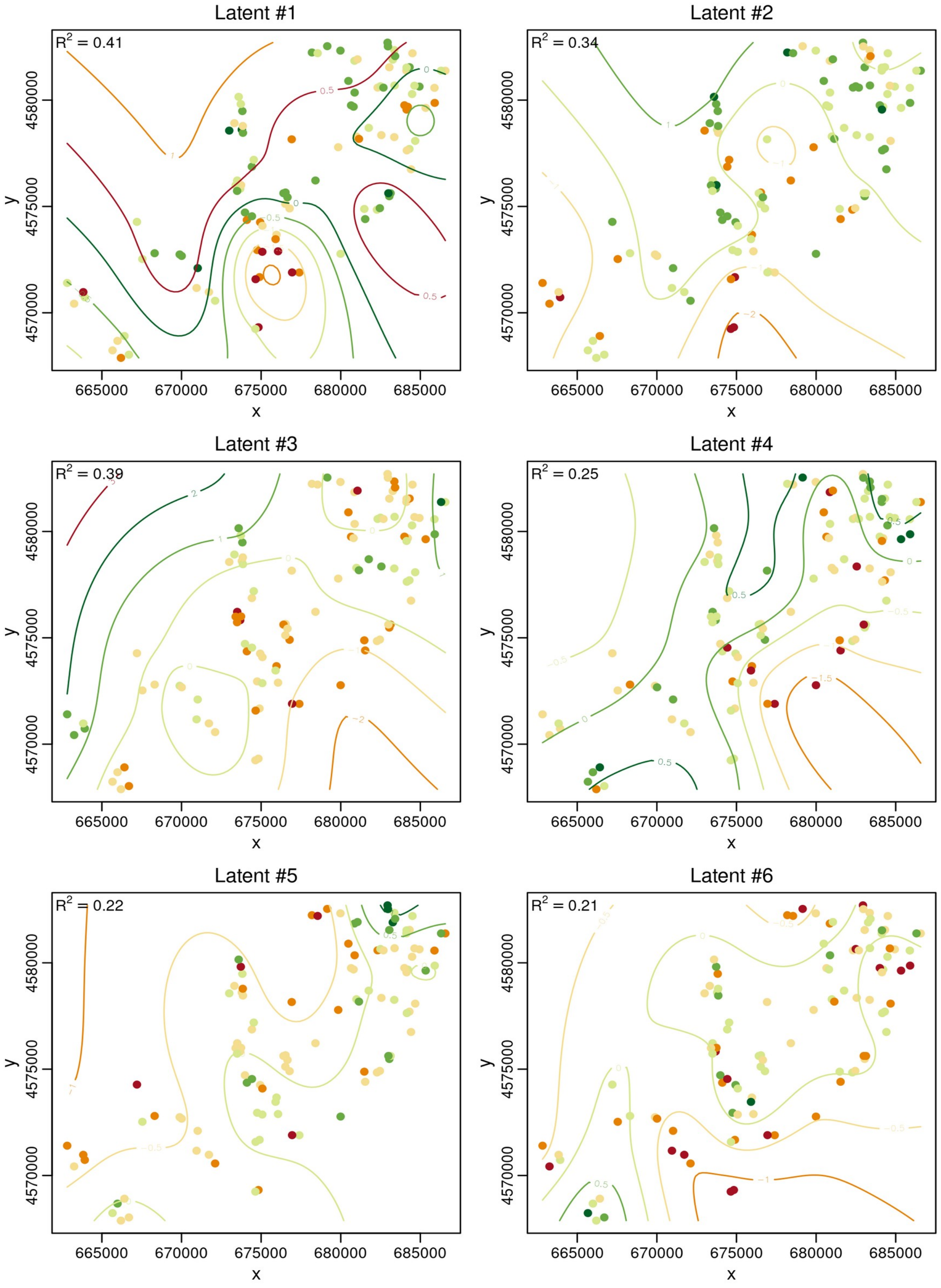

**Figure S2.** Model quality assessment (accuracy, specificity, sensitivity and precision) of the SCEM model in respect to the identified species interactions, for simulated communities with 64 species generated with 50 true species interactions, under different hyperparameterizations (x axis). The x axis represents a combination of the penalty applied to the interaction coefficients $\lambda_\theta$ and the penalty applied to environmental coefficients $\lambda_\beta$. For accuracy and sensitivity, we only counted undetected interactions whose absolute true value was higher than 0.5 as false negatives, thereby giving focus to the most serious false negatives (i.e. strong interactions that were not detected). For all calculations, we focused on the detection of interactions regardless of their direction, since the estimation accuracy of direction was analysed separately (Fig. S4). See main text and Appendix S3 for details.

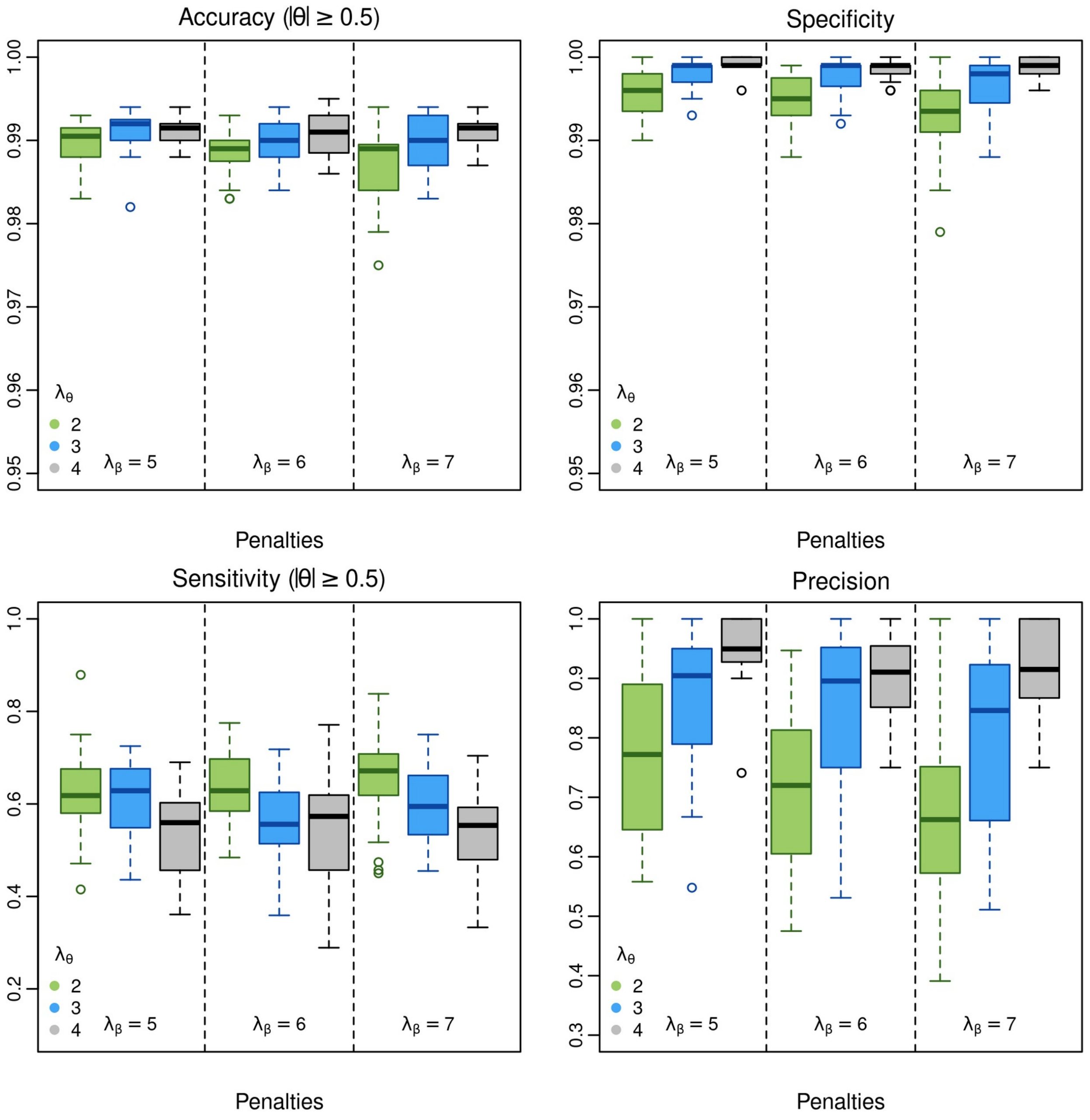

**Figure S3.** Degree of match between the estimated and true coefficients of (a) detected interactions (disregarding direction) and (b) environmental responses (first latent variable) of virtual communities with 64 species-50 interactions. Both are measured by the $R^2$ of a linear model between the estimated and the true values. In (b), the linear model models the first estimated latent variable values as a function of the true values of all predictors, because of rotational invariance of the latent variables. See main text and Appendix S3 for details.

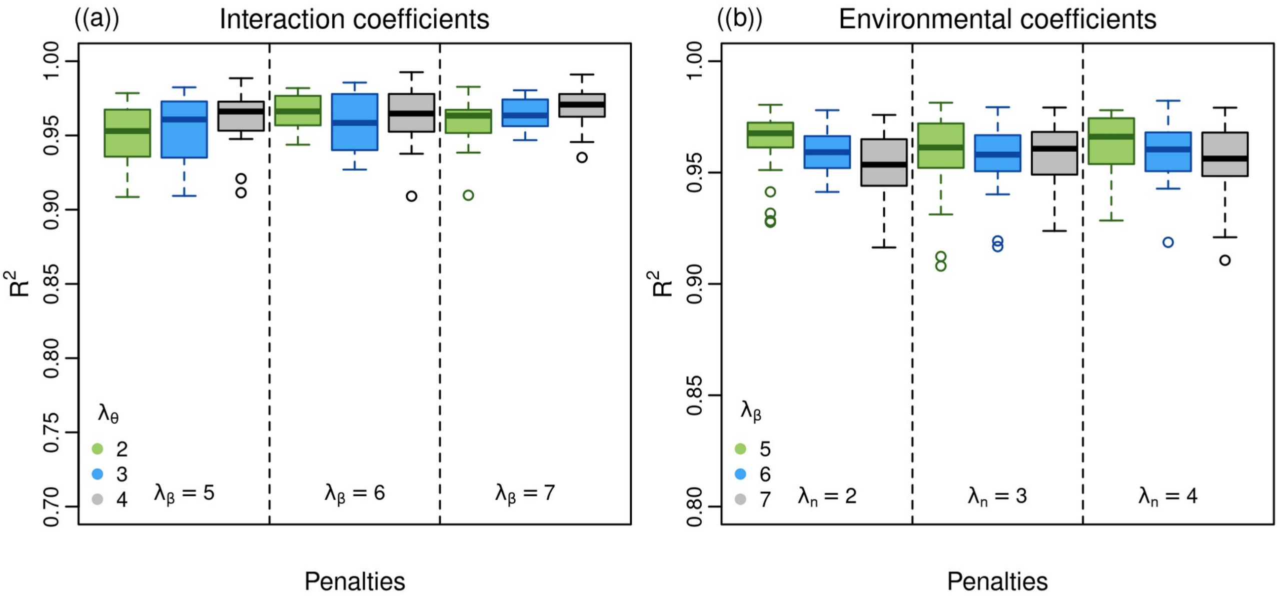

**Figure S4.** Accuracy of the SCEM model in recovering the true direction of the detected interactions (true positives, TP), for virtual communities with 64 species-50 interactions, under different hyperparameterizations (x axis). Note that interaction direction (i.e. species A affects species B vs. B affects A) is not related with sign (i.e. the type of interaction, negative or positive).

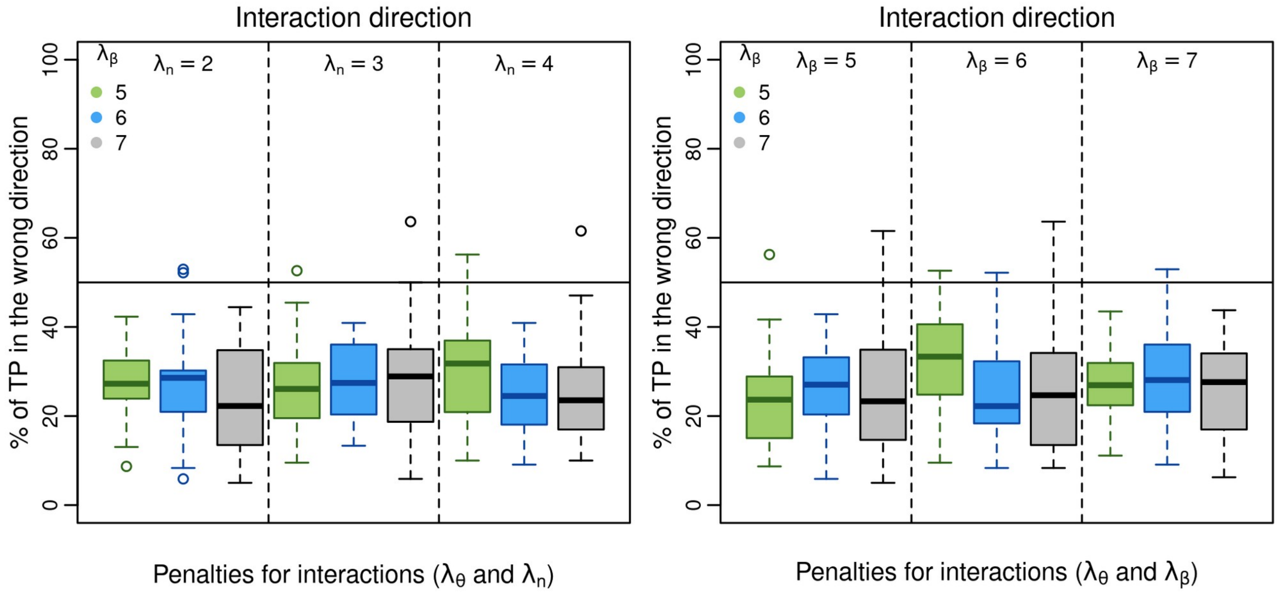

**Figure S5.** Estimated latent variables of a SCEM plotted against the true environmental predictors, for two fitted models with 64 species-50 interactions (top) and 128 species-100 interactions (bottom). The true predictors were dropped for model fitting to assess the ability of the model to recover their values as latent variables. Latent variables were standardized after estimation. Due to rotational invariance of the latent variables, for the purposes of this plot only, the rotation maximizing the correlation of each latent variable with each of the true predictors is shown.

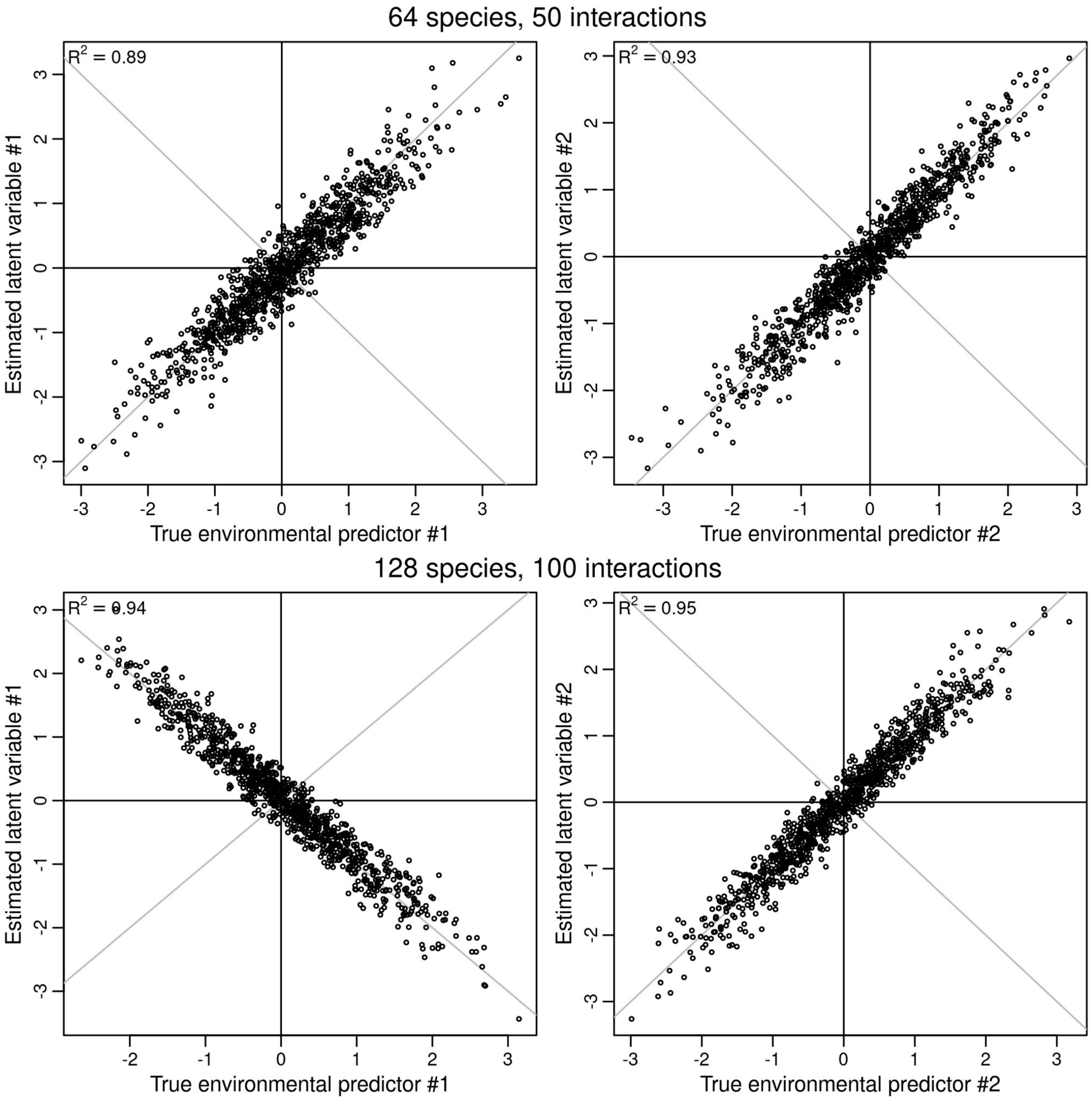

**Figure S6.** Estimated species interactions versus true values of an estimated SCEM model (on a virtual community with 128 species-100 interactions), before network selection, i.e. under a full network, where all possible interactions in both directions (n=16 256) are included and estimated. Note that, despite the large amount of false positives (red circles), their coefficients are small due to LASSO regularization, and there is a good match between the estimates of the true positives and their true values, although a small bias due to penalization is visible at the extremes. For each interaction, only the direction with the highest coefficient (in absolute value) is plotted. Green circles represent those whose highest coefficient (in absolute value) corresponds to the true direction, blue circles otherwise.

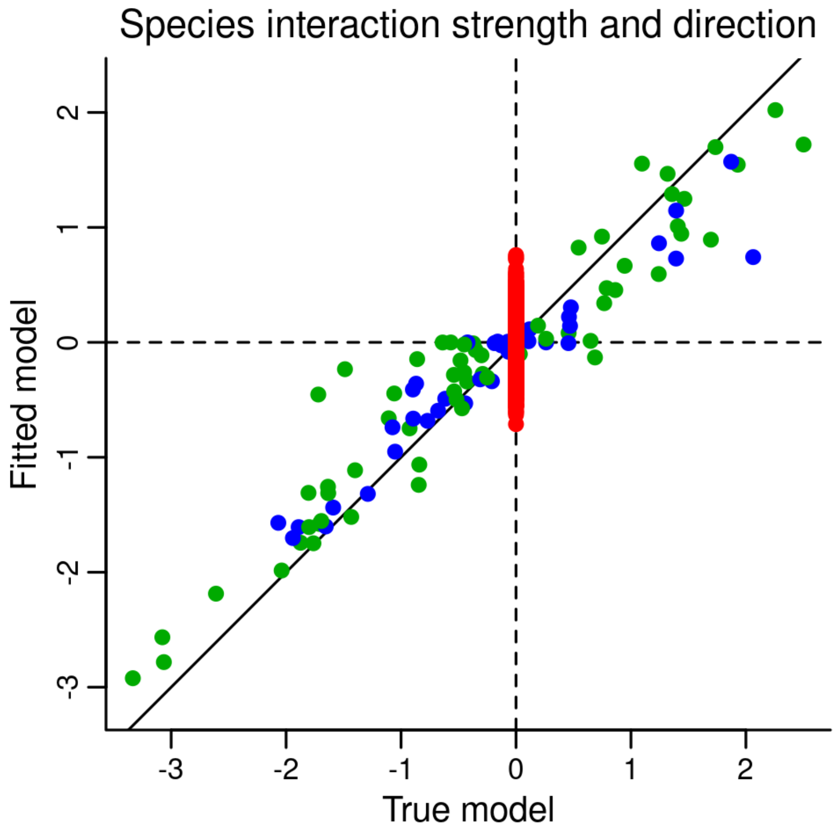

**Figure S7.** Estimated latent variables of the SCEM model fitted to the real occurrence data (y axis) plotted against the site scores of a partial Principal Component Analysis of the same data, removing the effects of the six environmental variables (x axis). Axes are both centered and scaled to unit variance. Each latent factor was matched with the PCA axis that yielded the highest Pearson correlation.

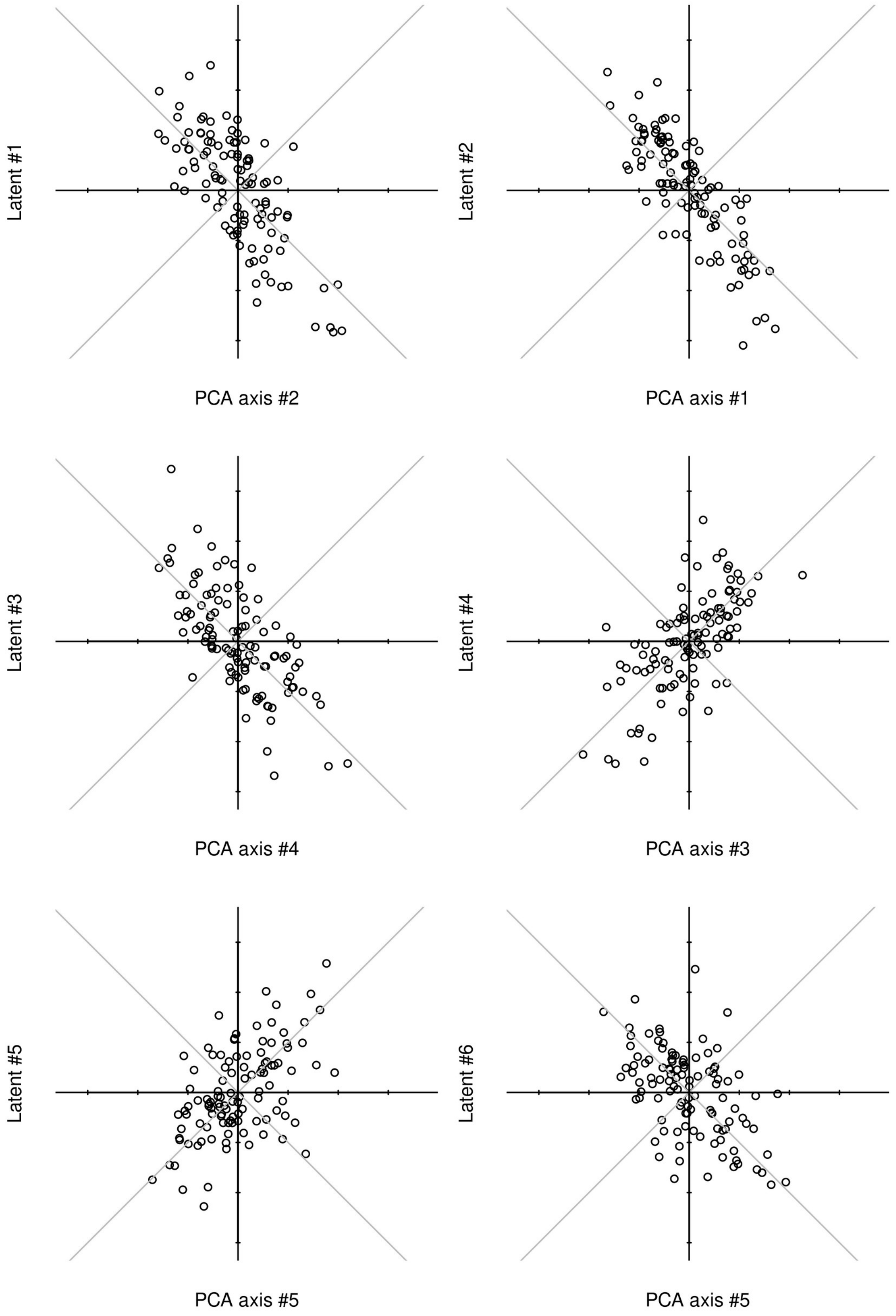
